# Charting DENR-dependent translation reinitiation uncovers predictive uORF features and links to circadian timekeeping via Clock

**DOI:** 10.1101/407551

**Authors:** Violeta Castelo-Szekely, Mara De Matos, Marina Tusup, Steve Pascolo, Jernej Ule, David Gatfield

**Affiliations:** Center for Integrative Genomics, University of Lausanne, Genopode, 1015 Lausanne, Switzerland.; Department of Dermatology, University Hospital of Zurich, Gloriastrasse 31, 8091 Zurich, Switzerland.; Faculty of Medicine, University of Zurich, 8091 Zurich, Switzerland.; Department of Neuromuscular Diseases, UCL Institute of Neurology, Queen Square, London WC1N 3BG, UK.; The Francis Crick Institute, 1 Midland Road, London NW1 1AT, UK.

## Abstract

The non-canonical initiation factor DENR promotes translation reinitiation on mRNAs harbouring upstream open reading frames (uORFs). Moreover, DENR depletion shortens circadian period in mouse fibroblasts, suggesting involvement of uORF usage and reinitiation in clock regulation. To identify DENR-regulated translation events transcriptome-wide and, in particular, specific core clock transcripts affected by this mechanism, we have used ribosome profiling in DENR-deficient NIH3T3 cells. We uncovered 240 transcripts with altered translation rate, and used linear regression analysis to extract 5’ UTR features predictive of DENR dependence. Among core clock genes, we identified Clock as a DENR target. Using Clock 5’ UTR mutants, we mapped the specific uORF through which DENR acts to regulate CLOCK protein biosynthesis. Notably, these experiments revealed an alternative downstream start codon, likely representing the bona fide CLOCK N-terminus. Our findings provide insights into uORF-mediated translational regulation that can regulate the mammalian circadian clock and gene expression at large.

## INTRODUCTION

Initiation is considered the rate-limiting step for eukaryotic translation (1) and can be regulated by multiple mechanisms including through upstream open reading frames (uORFs), which are short translated sequences defined by a start codon within the 5’ untranslated region (5’ UTR) of the transcript and an in-frame stop codon. Canonical translation initiation involves association of the small ribosomal subunit with an mRNA’s 5’ cap structure and scanning of the 5’ UTR until a start codon is recognised. In cases when this translation event is on a uORF, the ribosome can resume scanning after termination, allowing reinitiation at the downstream protein coding sequence (CDS). Alternatively, a scanning ribosome might skip certain uORFs, in a process known as leaky scanning. Translation reinitiation and leaky scanning are poorly understood mechanisms that may involve canonical and non-canonical initiation factors and that are likely regulated by specific features of the uORF and within the transcript (reviewed in (2, 3)).

Density regulated protein (DENR) and Malignant T-cell amplified sequence 1 (MCTS1) form a heterodimeric non-canonical initiation factor that has been described to bind to the small ribosomal subunit and to promote the eIF2-independent recruitment of the initiator tRNA to viral mRNAs, as well as to participate in the recycling of posttermination 40S subunits (4, 5). In *Drosophila*, DENR-MCTS1 was identified as a reinitiation factor required for efficient CDS translation on transcripts containing short and strong uORFs (6). Furthermore, based on reporter assays with synthetic constructs containing uORFs of varying lengths and Kozak strengths, endogenous DENR targets have recently been predicted in human cells (7). However, an unbiased identification of DENR targets *in vivo* and a transcriptome-wide quantification of the effects exerted by DENR on translational regulation are still lacking.

uORF usage is pervasive in the mammalian transcriptome, with up to 40% of transcripts predicted to contain at least one uORF (8). It generally reduces CDS translation efficiency (TE) because scanning ribosomes are diverted from the main ORF and unlikely all reinitiate (3). We have previously reported that several transcripts encoding components of the circadian clock machinery contained translated uORFs (9). Moreover, DENR loss-of-function – as a means of globally interfering with uORF regulation – shortened the period of free-running circadian oscillations in mouse fibroblasts by up to 1.5-hours (9). However, it has remained elusive on which circadian clock transcripts and through which uORFs DENR can act.

Here, we have used ribosome profiling in mouse NIH3T3 fibroblasts to identify and quantify DENR-mediated translational regulation transcriptome-wide, and in particular within core clock genes. We have used comprehensive regression analysis to characterise 5’ UTR features associated with DENR-dependent translational regulation and validate targets identified from our translatome-wide approaches. Finally, we show that DENR regulates efficient CLOCK biosynthesis involving a strong uORF and a so-far unannotated CDS initiation codon.

## MATERIALS AND METHODS

### Cell culture

NIH3T3 and HEK293FT cells were cultured under standard conditions (DMEM; 10% FCS, 1% penicillin/streptomycin, all from Invitrogen; 37°C; 5% CO_2_).

Lentiviral particle production in HEK293FT cells using envelope pMD2.G and packaging psPAX2 plasmids, and viral transduction of NIH3T3 cells, were performed following published protocols (10), with puromycin selection at 5 *μ*g/ml for 4 days.

### Cloning and plasmids

For the generation of lentiviral shRNA expression vectors, two different sequences targeting *Denr* were cloned into pLKO.1puro backbone vector (Addgene no. 10878 (11)): shRNA1: GTACCACAGAAGGTCACGATA, corresponding to clone TRCN0000308443 of the TRC shRNA Library from the Broad Institute; and shRNA2: GTGCCAAGTTAGATGCGGATT, corresponding to clone TRCN0000098826. pLKO.1puro constructs containing *Gfp*, and Scramble (Addgene no. 1864) shRNAs served as controls.

For the generation of dual luciferase (Firefly/Renilla) reporter plasmids, fragments containing the 5’ UTR and first 5-14 codons of the selected DENR targets/controls were amplified by PCR from genomic DNA or cDNA, cloned into the BamHI site of the prLV1 dual luciferase reporter plasmid (12), and validated by sequencing. The following primers were used for PCR: *Etaa1*, forward (for): aaaggatccGACTTGCAA AGATGGCGCTGCAC, reverse (rev): tttggatccactagtGTCCT TCAGCTGCATTCACATTT; *Map2k5*, for: aaaggatccGGTT CCGGAGTAACAGCGGTCTAAC, rev: tttggatccactagtGGC CAGCCACAGCATTACAGGTTAA; *Lrrc28*, for: aaaggatcc GCCGCTGAGTGCCGGTCAGCGGGC, rev: tttggatccactag tGATTTCGGATGCCATGACTGAACA; *Klhdc8a*, for: aaag gatccGTCTCCGACCCTGTAGACACTGCAG, rev: tttggatc cactagtATTGGGCACTTCCATGGCAGCCCGG; *Clock*, for: aaaggatccGGGGAGGAGCGCGGCGGTAGCGGTG, rev: tt tggatccactagtAACAATTGAGCTCATTTTACTACAGCTT ACGG; *Fth1*, for: aaaaggatcCAGACGTTCTCGCCCAGAGTCGC, rev: tttggatcccatGGTGGCGGCGGGGCGAGGCGC; *Rpl30*, for: aaggatcCAGCGGCGGGTGGACAGAGATGCC, rev: tttggatccacgCGTCTTCTTTGCGGCCACCATCTTC; *Deaf1*, for: aaggatccGGGCGTCAGACTGCCTCTTCGGTC, rev: tttggatccacgcgtGGCAGCGGCCGCCACAGCTGCCGC; *Cry1*, for: aaaaagatctactagtCACCCGGGCAGCCTCGGGACCG, rev: tttagatctTCGGAACCAGTGCACGGCGTTCAC.

5’ UTR mutants were generated by site-directed mutagenesis of the corresponding plasmids, using the following primers (mutated nucleotides in small case with mutated start/stop codon underlined in the forward sequence): *Klhdc8a*-Mut, forward (for): GGCAGGGAGCCCGTGCAG CGCGgt<u>acc</u>GAGGCTGAGAGAGGGGACGCGCC, reverse (rev): GGCGCGTCCCCTCTCTCAGCCTCggtacCGCGCTGCACGGGCTCCCTGCC; *Map2k5*-Mut, for: CTCTTAC TCACAAGGACTACAG<u>gcta</u>gcTTGTGGGTTCTGTTTTG TCCAC, rev: GTGGACAAAACAGAACCCACAAgctagcCTGTAGTCCTTGTGAGTAAGAG; *Etaa1*-Mut, for: cga aggatcCGACTTGCAAAG<u>gta</u>ccGCTGCACGGTGGCGC GCGGGCTC, rev: GAGCCCGCGCGCCACCGTGCAGC ggtacCTTTGCAAGTCGgatccttcg; *Clock*-uORF-Stop-Mut, for: GTTATGGTGTTTACCG<u>TcA</u>GCTGTAGTAAAAT GAG, rev: CTCATTTTACTACAGCTgACGGTAAACACCATAAC; *Klhdc8a*-uORF-Stop-Mut, for: GCCGACCG GCGCCGTG<u>cac</u>CCCGCGGGCCCCGGGGCT, rev: AGC CCCGGGGCCCGCGGGgtgCACGGCGCCGGTCGGC; *Etaa1*-uORF-Stop-Mut (small case correspond to an insertion that changes uORF frame), for:TGCTCCGCTGTAGCAgc AAATGTGAATGCAGC, rev: GCTGCATTCACATTTgcT GCTACAGCGGAGCA; Clock-uORF1-Mut, for: CCCAAA ATCACCAGCAAGAGTTgctagcGTCAGTCACACAGAA GACGGCC, rev: GGCCGTCTTCTGTGTGACTGACgctagcAACTCTTGCTGGTGATTTTGGG; *Clock*-uORF2-Mut, for: AAAGTGAAAGAGGAGAAGTACAggtacCTACCACAAGACGAAAACATAAT, rev: ATTATGTTTTCGT CTTGTGGTAGgtaccTGTACTTCTCCTCTTTCACTTT; *Clock*-AnnotCDS-Mut, for: CAAGACGAAAACATAATGT GTT<u>tcG</u>GTGTTTACCGTAAGCTGTAG, rev: CTACAGC TTACGGTAAACACCgaAACACATTATGTTTTCGTCTT G; *Clock*-AlterCDS-Mut, for: GTTTACCGTAAGCTGTAG TAAA<u>caG</u>AGCTCAATTGTTactagtg, rev: cactagtAACAAT TGAGCTCtgTTTACTACAGCTTACGGTAAAC.

### iCLIP experiments and analyses

For iCLIP experiments, we either used NIH3T3 cells without overexpression to immunoprecipitate the endogenous DENR, or NIH3T3 cells transiently transfected with *Denr* cloned into a FLAG-Venus-pLenti6 vector (Addgene 36948). Cells were grown to confluency in a 15-cm dish. One dish was used for overexpression, and six for non-transfected experiments.

iCLIP was performed as described (13, 14). RNA-protein complexes were crosslinked with UV-C (254 nm) at 150 mJ/cm^2^ in a Stratalinker 2400. After cell lysis (50 mM Tris-HCl, pH 7.4; 100 mM NaCl, 1% Igepal (Sigma), 0.1% SDS, 0.5% Na deoxycholate, EDTA-free protease inhibitors), RNA was partially digested with 0.2 U RNase I (Thermo Fisher). DENR-RNA complexes were pulled down with anti-DENR antibody (Abcam 108221), pre-immobilised on protein G-coated magnetic beads (Dynal). An IgG immunoprecipitation was used as control. Immuno-complexes were washed twice with High Salt Buffer (50 mM Tris-HCl pH 7.4, 1 M NaCl, 1 mM EDTA, 1% Igepal, 0.1% SDS, 0.5% sodium deoxycholate) and twice with PNK Wash Buffer (20 mM Tris-HCl pH 7.4, 10 mM MgCl^2^, 0.2% Tween-20). An infrared adaptor was ligated to RNA 3’ ends, and protein-RNA complexes were visualised in a LiCor CLx Imager (Odyssey), after denaturing PAGE and nitrocellulose membrane transfer. 60-130 kDa DENR-RNA complexes were excised from the membrane and liberated by proteinase K (Roche) digestion. This size range likely reflects DENR-MCTS1 complexes (26 kDa and 22 kDa, respectively) bound to RNA, rather than DENR-RNA alone, indicating that the DENR-MCTS1 interaction is strong. During library preparation, cDNAs were ligated on their 5’ end to a 5-nt sample barcode to allow demultiplexing. The barcode is preceded by a 4-nt and followed by a 3-nt random sequence that serves as unique molecular identifier (UMI) to mark individual cDNA molecules. cDNA libraries were then PCR amplified (24 cycles) and subjected to high-throughput sequencing (Illumina HiSeq2500 platform). Experiments were done in triplicate for DENR-overexpression, and in duplicate for non-transfected and IgG IP conditions.

iCLIP data was processed and analysed using the iCount platform (https://github.com/tomazc/iCount), as described (15). Briefly, the sample barcodes were removed, and the UMI was excised and added to the read ID prior to mapping with Bowtie 0.12.7 to the mouse genome (mm10), allowing two mismatches. Sequences with the same UMI and starting at the same genomic position – likely representing PCR duplicates – were excluded from subsequent analyses. High-confidence crosslink sites were calculated with the iCount function *peaks*, with options *RegionsAsOne* and 15-nt flank.

### Ribosome profiling

One 15-cm dish of confluent NIH3T3 cells was used per replicate. Ribosome profiling and parallel RNA-seq were performed in triplicate for *Denr* shRNA1, *Denr* shRNA2 and scr shRNA, and in duplicate for *Gfp* shRNA, following the ART-Seq protocol by Epicentre-Illumina (Cat.no. RPHMR12126) with minor modifications as described below. Cells were treated with cycloheximide (100 *μ*g/ml) for 2 min at 37°C to arrest translation elongation, and immediately harvested, followed by lysate preparation. For the generation of ribosome-protected fragments, cell lysates were treated with 5 units RNase I (ART-Seq) per OD260 and monosomes were purified in MicroSpin S-400 columns (GE Healthcare). For both RPF- and RNA-seq, 5 *μ*g of RNA were used for rRNA depletion following the Ribo-Zero Magnetic Kit instructions from Epicentre-Illumina (Cat.no. MRZH11124). Finally, libraries were amplified using 12 PCR cycles, and sequenced (HiSeq2500 platform).

### Sequencing pre-processing, alignment and quantification

Initial processing, mapping and quantification of mRNA and footprint abundance were performed as previously (16). Briefly, adaptor sequence was trimmed with Cutadapt (17) and trimmed sequences were filtered using a custom python script to keep 26-35 nt and 21-60 nt reads for RPF- and RNA-seq, respectively. Reads were then sequentially aligned to mouse rRNA, tRNA and cDNA (Ensembl mouse release 84) using Bowtie2 (18) and mouse genome using Tophat v2.0.11 (19). Trimmed and filtered sequences from RNA-seq were also directly mapped against the mouse genome with Tophat to estimate expressed transcript isoforms (using Cufflinks v2.2.1 (20)). Transcriptome-mapping reads from the sequential alignment were counted towards their location on 5’ UTR, CDS or 3’ UTR per gene, using only the set of expressed isoforms as estimated above. The location of the putative A-site of the footprint was used for RPF-seq counting (position +15 from the 5’ for reads ≤30 nt and position +16 for reads >30 nt), and the 5’ end was used for RNA-seq reads. For genes with multiple expressed isoforms, reads that did not map unambiguously to a single feature were assigned to one in preference order: CDS, 5’ UTR, 3’ UTR.

RPF- and RNA-seq read counts were normalized separately with the upper quartile normalization from edgeR (21) and then transformed to RPKM values as read count per gene kilobase per normalized library size. Genes with less than 10 reads in 2/3 of samples and mitochondrially-encoded genes were removed before normalization.

Translation efficiencies were then calculated per sample as the ratio of RPF RPKM to RNA RPKM of the region of interest (5’ UTR, CDS or uORF), log2 transformed and averaged over all replicates.

### Differential translation efficiency analysis

Significant changes in TE between knockdown and control cells were assessed with several available algorithms: Xtail (22) with options *bins=10000*, RiboDiff (23) with options - *d 1 -p 1*, Riborex (24) with options *engine=“DESeq2”*, and Babel (25) with options *nreps=100000*.

Briefly, all four methods, with the options implemented in our study for maximal consistency among them, model raw read counts by following a negative binomial distribution, and use global normalization methods for sequencing depth that would not account for global changes in TE if existing. RiboDiff and Riborex use DESeq2 to estimate a single mean and dispersion for both RPF- and RNA-seq, and detect differentially translated genes by fitting RPF- and RNA-seq read counts to a single GLM, whereas Xtail uses DESeq2 for mean and dispersion estimation separately for RPF- and RNA-seq and fits two independent GLMs. Xtail then derives probability distributions for fold changes of RNA or RPF, or for RPF-to-RNA ratios, and evaluates statistical significance between conditions from one of the two (the most conservative) pipelines. Babel relies on edgeR for parameter estimation and uses an errors-in-variables regression to test for significant changes in translation within and between conditions.

Differential TE was set at FDR < 0.1 for all four methods and performance for false positive discovery rate was further assessed by randomly label-shuffling knockdown and control datasets, which should not yield differential TE genes.

### uORF annotation and translation analyses

For uORF detection, a set of transcripts was used for which reads can be unambiguously assigned to the 5’ UTR, i.e. transcripts that are the only protein-coding isoform expressed (N = 4637). In order to include as many transcripts as possible for the annotation, genes with multiple protein-coding isoforms were also considered if 1) one isoform represented > 65% of all expressed (as derived from our transcript expression estimation, see “Sequencing preprocessing, alignment and quantification”, N = 3211) or 2) all expressed isoforms had the same CDS start, in which case the transcript with the highest expression was selected (N = 1826). For the selected transcripts, uORFs were annotated and considered as translated with the following criteria: 1) started with AUG, CUG, GUG or UUG, 2) had an in-frame stop codon within the 5’ UTR or overlapping/within the CDS, 3) were at least 9 nt long (including stop codon), 4) had a coverage > 25%, 5) showed a frame preference and 6) the preferred frame was the one corresponding to the uORF 5’ end/start codon. Importantly, we considered only uORFs coding for at least 2 amino acids (i.e. 9 nt, including the stop codon), as for shorter uORFs frame bias assessment would be highly prone to false positives, and Kozak context scoring would be compromised (for 6 nt uORFs, the position +4 would always be the T of the stop codon, and thus cannot be a G of the strongest Kozak context).

### Multiple regression analysis

For the modelling, all transcripts containing translated uORFs in *Denr* knockdown or control cells were included (N = 4755) and the following 5’ UTR features were used as predictors of the fold change in CDS TE across conditions: 1) the number of uORFs, 2) the uORFs length, using the median length in transcripts with several uORFs, 3) the proportion of AUG uORFs, 4) the uORF Kozak context, scored relative to the consensus GccA/Gcc###G, where upper-case letters denote highly conserved bases (scored +3), lower-case letters indicate the most common nucleotides (scored +1) and ### represents the start codon (not scored). The median Kozak score was used when several uORFs were present on a transcript. 5) the CDS Kozak context, scored in the same manner, 6) the distance of the first uORF start codon from the 5’ cap, 7) the distance of the last uORF stop codon to the CDS start, 8) the uORF GC content, using the median for transcripts with several uORFs, 9) the presence of overlapping uORFs in the transcript (binary predictor). Predictors with skewed distributions (number of uORFs, uORF length, distance from the cap and distance to the CDS start) were log2 transformed. Partial regression plots were generated with the R package ‘*visreg*’. Absence of multicollinearity between predictors was assessed with the variance inflation factor using R function *vif* (*car* package).

### Functional enrichment analysis

Functional enrichment analysis was performed using g:Profiler (26), including Gene Ontology Biological Process and Molecular Function terms and biological pathway information from Reactome. Functional categories of more than 2 and less than 1500 members were used and p-values were corrected by Benjamini-Hochberg FDR. Categories with FDR < 0.05 were considered as significant and were used for visualization with Cytoscape v.3.5.1 EnrichmentMap tool (27).

### Dual luciferase assay

NIH3T3 cells were first lentivirally transduced with the dual luciferase reporters containing the 5’ UTR construct of interest and then transduced with *Denr*-shRNA/scramble-shRNA. After puromycin selection, cells were lysed in passive lysis buffer and luciferase activities were quantified using DualGlo luciferase assay system and a GloMax 96 Microplate luminometer (all from Promega). Firefly/Renilla luciferase (FL/RL) activities were first normalized to that of the empty vector in scramble shRNA-transduced cells. For DENR target experiments, FL/RL signals were also normalized to the scramble shRNA-transduced, non-mutated construct. For *Clock* mutational experiments, FL/RL signals were secondly normalized to the scramble shRNA-transduced *Clock*-WT construct.

### In *vivo* bioluminescence imaging

mRNAs were produced using in vitro transcription at the “ivt mRNA production and formulation platform” in Zurich (http://www.cancer.uzh.ch/en/Research/mRNA-Platform.html). The 5’ end consisted of a CleanCapTM (Trilink) followed by the WT or mutated *Klhdc8a* 5’ UTR and by a codon optimised firefly luciferase open reading frame. The 3’ end consisted of a tandem repeat of the mouse beta-globin 3’ UTR and a poly-A tail (28). Uridine residues were fully substituted by 1-methyl pseudouridine residues (this modification can prevent immunostimulation, avoiding production of type I interferon and consecutive inhibition of translation *in vivo* (28)).

Animal procedures were approved by the Veterinary Office and its research ethics review committee of the University of Zurich (Kanton Zurich, Health Direction, Veterinary Office, Zollstrasse 20; 8090 Zurich; license number ZH215/17). Animals were purchased from Envigo (Netherlands). 4 to 8-week-old mice were injected intravenously with 1 *μ*g of mRNA formulated with TransIT transfection reagent following the manufacturer’s suggestions (Mirus). The procedure for one mouse was as follows: the nucleic acid was diluted in 38 *μ*l of Opti-MEM (Gibco) before the addition of 0.72 *μ*l of the “mRNA Boost Reagent”. After mixing, 1.12 *μ*l of the TransIT reagent were added. The solution was thoroughly homogenated and injected immediately: 40 *μ*l per mouse. At different time points after mRNA injection, *in vivo* bioluminescence imaging was performed on an IVIS Lumina instrument (PerkinElmer). Immediately before each measurement, D-luciferin (Synchem) dissolved in PBS (15 mg/ml stock) and sterile filtered was injected (150 *μ*g/g intraperitoneally). Emitted photons from live animals were quantified 10-20 min post luciferin injections, with an exposure time of 3 min. Regions of interest (ROI) were quantified for average radiance (photons s-1 cm-2 sr-1) (IVIS Living Image 3.2).

### Western blotting

Total protein extracts from NIH3T3 cells were prepared according to the NUN procedure (29). Antibodies used were rabbit anti-CLOCK 1:3000 (ab3517, abcam), mouse anti-U2AF65 1:5000 (Sigma), rabbit anti-DENR 1:4000 (ab108221, abcam), and anti-rabbit-HRP or anti-mouse-HRP 1:10000 (Promega). Blots were visualised with the Fusion Fx7 system and quantified with ImageJ 1.48k.

## RESULTS

### Redistribution of ribosomes from CDS to 5’ UTRs upon *Denr* knockdown

We hypothesized that if DENR is required for efficient protein production from the main CDS of mRNAs that undergo reinitiation after uORF translation, we would be able to identify DENR-regulated transcripts by comparing ribosome occupancies transcriptome-wide in DENR loss-of-function vs. control cells by ribosome profiling. Conceivably, DENR target mRNAs should exhibit decreased translation efficiencies (TEs) on their CDS when DENR is absent (Fig. 1A). We therefore performed shRNA-mediated knockdown of *Denr* in mouse NIH3T3 fibroblasts, a cell line in which we had previously observed that *Denr* depletion led to a shortening of the period of circadian oscillations (9). Cells were lentivirally transduced with two different *Denr*-targeting shRNAs (shRNA1, shRNA2; in triplicate) or control RNAs (scramble shRNA, in triplicate; *Gfp* shRNA in duplicate), grown to confluency and harvested in the presence of cycloheximide (CHX) to stabilize elongating ribosomes. After preparation and sequencing of ribosome-protected mRNA fragment (RPF-seq) and total RNA libraries (RNA-seq) as previously described (9), we obtained ribosome footprints that were predominantly 29-30 nt in length (Supplementary Fig. 1A), that preferentially mapped to protein coding sequences (Supplementary Fig. 1B) and that showed the characteristic triplet periodicity expected from high-quality footprint data (Supplementary Fig. 1C). Across replicates, CDS-mapping normalised reads were highly correlated (R^2^ > 0.97, Supplementary Fig. 1D), indicating good reproducibility. The RNA-seq and RPF-seq data confirmed the efficiency of *Denr* knockdown (77-90% reduction in *Denr* transcript levels), whereas the abundance of *Mcts1* mRNA, encoding the DENR heterodimerization partner, or of *Eif2d*, encoding a structurally and functionally related homologue of the DENR-MCTS1 dimer (4), remained unchanged (Fig. 1B, C and Supplementary Fig. 1E).

**Figure 1.**
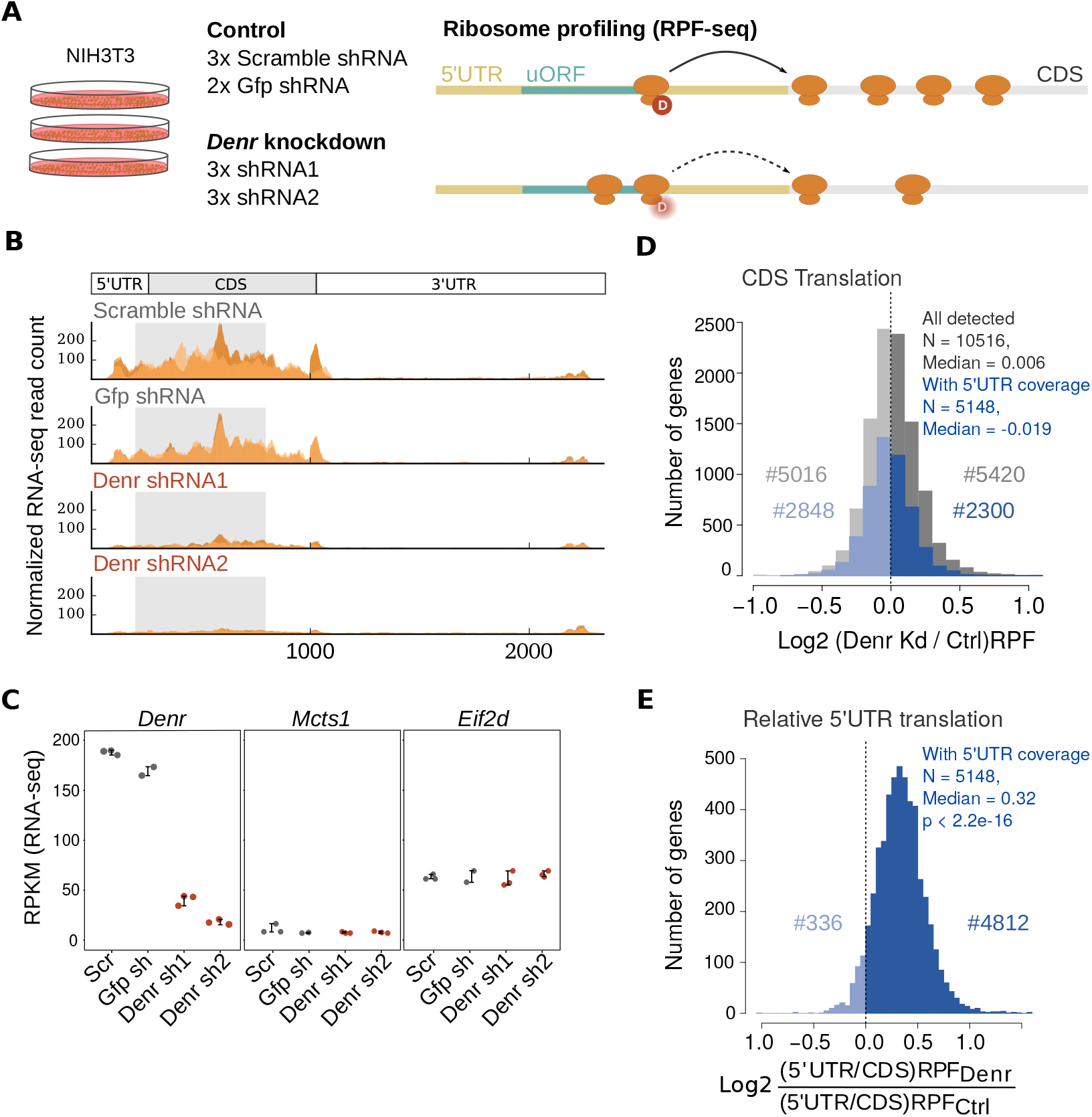
*Denr* knockdown engenders ribosome redistribution from CDS to 5’ UTR. **A)** Experimental approach of the study. Ribosome profiling (RPF-seq) and parallel RNA-seq were performed on NIH3T3 mouse fibroblasts depleted of *Denr* via lentivirally expressed shRNAs, with scramble (Scr) and *Gfp* shRNAs as control conditions. Profiling was carried out in biological triplicates for the two *Denr* shRNAs (shRNA1, shRNA2) and for Scr shRNA, and in duplicate for *Gfp* shRNA. We hypothesized that if DENR is required for efficient reinitiation after uORF translation, knocking down *Denr* would lead to reduced translation efficiency on the main coding sequence of DENR targets. **B)** Distribution of normalized RNA-seq read counts along the *Denr* transcript show high *Denr* knockdown efficiency. Different orange color shadings correspond to the different replicates. **C)** RPKM values for CDS-mapping RNA-seq read counts for *Denr* (left), *Mcts1* (middle), and *Eif2d* (right) indicate specific and high knockdown efficiency. Each dot corresponds to a sample and replicate and error bars indicate mean±sd. **D)** Distribution of CDS translation (from RPF-seq) differences between *Denr* knockdown (Kd) and control cells. In grey, all detected genes (N = 10516); in blue, genes with sufficient coverage in 5’ UTRs (i.e. a minimum of 20 RPF-seq reads in the 5’ UTR in 3/4 of samples; N = 5148). Note that both distributions are centered around zero, indicating that *Denr* knockdown does not strongly affect global protein synthesis *per se*. **E)** Distribution of 5’ UTR-to-CDS translation differences between *Denr* Kd and control cells, for genes with sufficient 5’ UTR coverage (N = 5148). Relative 5’ UTR translation was significantly higher in *Denr* Kd cells (median = 0.32, p < 2.2e-16, Wilcoxon signed rank test), suggesting a ribosome footprint redistribution to 5’ UTRs upon *Denr* knockdown.

A reduction in *Denr* levels that alters reinitiation efficiencies should engender differences in ribosome occupancies on the CDS and, possibly, on uORFs. We first compared CDS footprint levels for all expressed genes (N = 10516) and noted no major global effects of *Denr* knockdown (Fig. 1D, grey). We then selected the subset of transcripts that also displayed footprint coverage within the 5’ UTR (N = 5148). Of note, the quantification of footprints from the full 5’ UTR can serve as a useful proxy for uORF translation (30, 31), given that uORFs are considered the major source of 5’ UTR-mapping reads. This group of genes showed an overall similar distribution as all genes (Fig. 1D), albeit with a slight shift to lower levels of CDS-mapping footprints upon *Denr* knockdown (p < 2.2e-16, K-S test). We concluded that also for this subset of genes, the global effect on translational output was relatively small. We next calculated the ratio of 5’ UTR-mapping to CDS-mapping footprints per gene, and determined how this measure of ‘‘relative 5’ UTR translation” was affected in *Denr*-depleted vs. control cells. Our analysis revealed a striking increase in relative translation on 5’ UTRs in *Denr*-deficient cells, indicative of a redistribution of translating ribosomes within transcripts (Fig. 1E and Supplementary Fig. 1F). This effect was absent from the RNA-seq data (Supplementary Fig. 1G). Conceivably, these changes on the translational landscape may reflect complex alterations in initiation efficiency and uORF usage, and are in line with a role of DENR in translation reinitiation (6).

Translational remodelling that affects a sizable proportion of the transcriptome is most consistent with a scenario in which DENR interacts with ribosomes at mRNAs with little, or no, sequence specificity. To address this question, we examined the direct interactions of DENR with RNAs *in vivo* by performing individual nucleotide resolution crosslinking and immunoprecipitation (iCLIP) (15) of DENR (Supplementary Fig. 2A). Over 85% of iCLIP reads mapped to ribosomal RNA, in particular to 18S rRNA (Supplementary Fig. 2B, C), with the five most prominent binding sites on helices 24-26a (Supplementary Fig. 2D, E). This outcome is in line with structural data that places the DENR-MCTS1 complex in proximity to the mRNA channel within the small ribosomal subunit and that identifies helix 24 as a potential interface of MCTS1 with 18S rRNA (32, 33). Moreover, helix 26 has previously been reported to make contacts with sequences upstream of restarting AUG codons on viral RNAs (34). In contrast, only 10.5% of iCLIP reads mapped to mRNAs, which was not sufficient to identify specific mRNA binding sites. In summary, while we cannot exclude transient direct interactions of DENR-MCTS1 with mRNAs, our iCLIP data are consistent with structural studies and support a model in which the complex binds close to the mRNA channel of initiating ribosomes at a location that could facilitate reinitiation when uORFs are present, but travels indiscriminately along mRNAs.

### Decreased CDS translation efficiency, uORF enrichment, and pathway analysis of DENR-regulated transcripts

From the shift in the distribution of 5’ UTR vs. CDS footprints that was affecting a significant proportion of the transcriptome in *Denr*-deficient cells (Fig. 1E), we wished to narrow down to the most-DENR-sensitive transcripts that were potentially direct and functionally relevant targets. We reasoned that changes in CDS translation efficiency (TE; defined as the ratio of CDS-mapping RPF-seq to RNA-seq reads) would represent an appropriate measure to identify such targets, as TE differences would alter the amount of protein produced from the transcript. We therefore calculated transcriptome-wide TEs and, given that there is currently no established consensus on the most suitable analytical tool for differential translation analysis, we then applied several available algorithms to detect significant TE changes. Briefly, the four methods we employed, Xtail (22), RiboDiff (23), Riborex (24), and Babel (25), are all conceived for the quantification of translational changes that deviate from possible variations already occurring at the level of mRNA abundances. They differ in parameter estimation and modeling approaches, as well as in sensitivity, false discovery rates and computational speed. When applied using settings that maximize comparability of results (see Materials and Methods), we identified 240, 78, 41 and 30 genes that had a significant change in TE between *Denr*-deficient and control cells with Xtail, RiboDiff, Riborex, and Babel, respectively (FDR < 0.1 for all four algorithms). Importantly, differential TE genes detected by Riborex, RiboDiff and Xtail completely overlapped, while only about half of the genes detected by Babel were also found by any of the other methods (Supplementary Fig. 3A). Moreover, most genes detected by Xtail, RiboDiff and Riborex showed decreased TE in *Denr*-depleted cells, as expected for the knockdown of a reinitiation factor, and they spanned the full range of expression levels (Fig. 2A, and Supplementary Fig. 3B, C). By contrast, Babel was biased towards low abundance transcripts and identified similar numbers of up- and down-regulated TE genes (Supplementary Fig. 3B, C). We concluded that Xtail likely represented the most sensitive method to identify DENR targets, and further estimated the false discovery rate of the algorithm by randomly shuffling the dataset labels, which eliminated nearly all differential TE genes (Supplementary Fig. 3D). Please note that, for simplicity, we shall further use the term “DENR targets” for the 240 differential TE genes identified by Xtail, although in the absence of information on direct DENR-mRNA interactions, the term “*most DENR-sensitive transcripts*” would be more precise.

**Figure 2.**
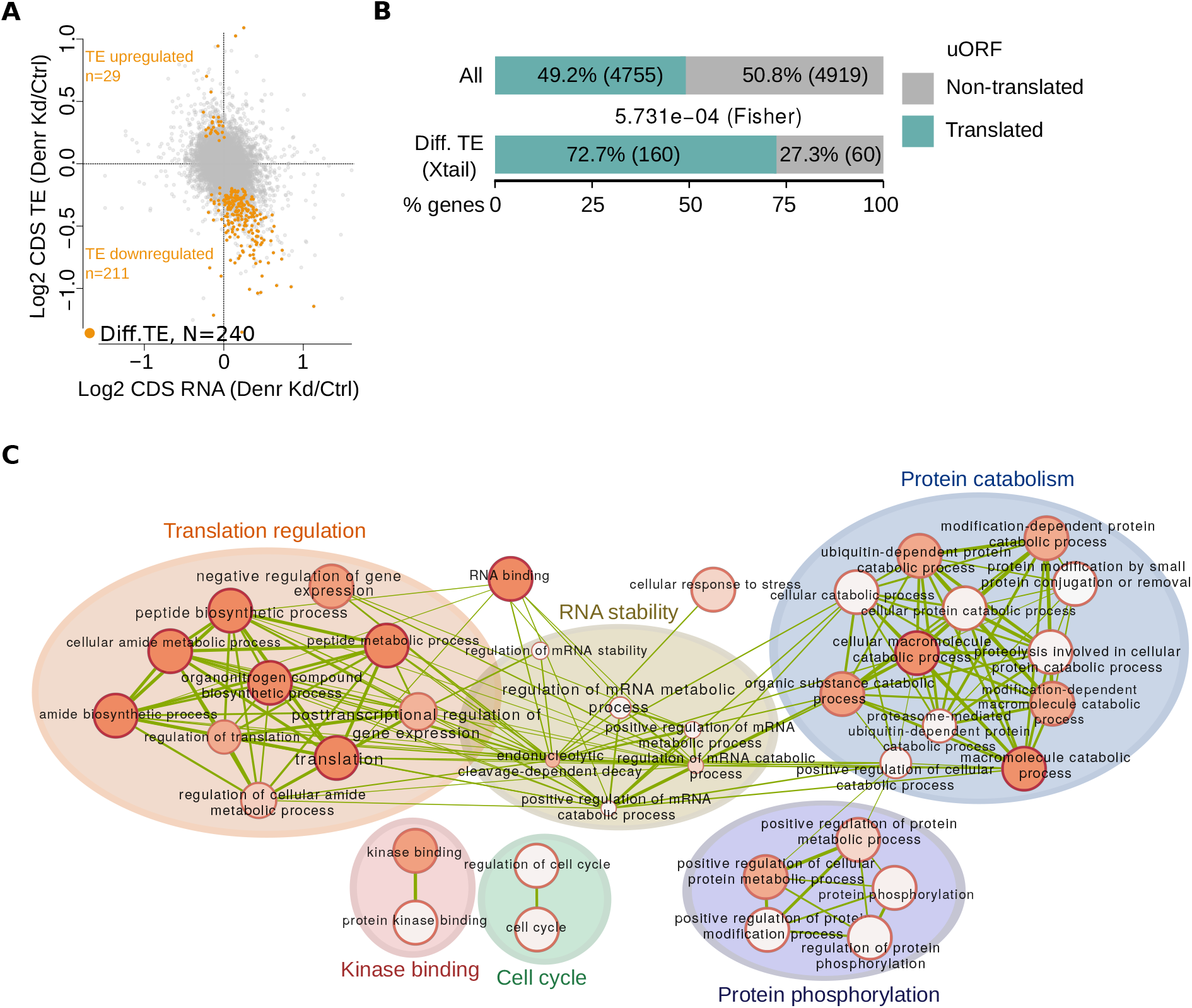
Characteristics of DENR targets: decreased CDS TE; enrichment for uORFs; association with functional categories. **A)** Scatter plot of *Denr* Kd-to-control ratio of mRNA abundance vs. CDS translation efficiency (TE) for all expressed genes (N = 10516). Highlighted in orange are genes with differential TE (detected by Xtail (22), FDR < 0.1, N =240), the majority of which showed decreased TE in *Denr* Kd cells, in line with a role of DENR in regulating translation reinitiation. **B)** uORF enrichment in DENR targets. Proportion of genes with translated uORFs transcriptome-wide compared to that among the differential TE genes detected in (A). DENR targets showed enrichment for translated uORF-containing genes (p = 5.73e-4, Fisher test). **C)** Enrichment map for GO-Biological process, GO-Metabolic function and Reactome functional categories of genes with differential TE. Only categories with an FDR < 0.05 in the GO analysis were considered. Node size is proportional to the number of genes associated with the GO category, shading corresponds to the statistical significance, and edge width is proportional to the number of genes shared between functional categories.

Among the 240 differential TE genes identified by Xtail (see Supplementary file 1) were several for which the presence of regulatory uORFs has been described previously, such as *Atf5* (35), *Rnaseh1* (36, 37), *Jak2* and *Map2k6* (38). In order to investigate whether the TE decrease among DENR-sensitive transcripts was associated with uORF translation, we annotated uORFs transcriptome-wide. Briefly, we included AUG and non-canonical, near-AUG (i.e., CUG, GUG, UUG) initiation codons located in 5’ UTRs, and assessed the active translation of the corresponding ORFs by their footprint coverage, frame preference and triplet codon periodicity (Supplementary Fig. 4A). Using this approach, we detected translated uORFs in nearly half of all expressed genes (N = 4755 genes, Fig. 2B), consistent with previous predictions (8). For the majority of genes, the uORFs were fully contained in the 5’ UTRs (51.8%, N = 2464 genes), but genes with uORFs overlapping the CDS (19.5%, N = 926 genes), or that had both overlapping and non-overlapping uORFs (28.7%, N = 1365 genes) were frequent, too (Supplementary Fig. 4B). Notably, DENR targets were highly enriched for genes with translated uORFs (72.7%, p = 5.73e-4, Fisher test, Fig. 2B) that were non-overlapping uORFs (Supplementary Fig. 4B). The depletion of overlapping uORFs (Supplementary Fig. 4B) – indicating that they are insensitive to DENR regulation – is consistent with the idea that after terminating on an overlapping uORF, a ribosome would presumably not be able to backtrack to reinitiate on an upstream CDS start codon. Moreover, relative 5’ UTR translation in *Denr*-depleted vs. control cells calculated over the full 5’ UTR – as a proxy of uORF translation – was significantly higher for DENR targets than for the global set of uORF-containing genes (Supplementary Fig. 4C, D). In summary, these observations were fully consistent with an involvement of DENR in reinitiation and in efficient CDS translation of a subset of uORF-containing mRNAs.

We next analysed whether specific functional pathways were over-represented among the 240 differential TE genes. GO term analysis revealed enrichment of categories associated with translation and mRNA stability, as well as cellular functions related to protein phosphorylation and catabolism, kinase binding and cell cycle (Fig. 2C and Supplementary Fig. 4E), in line with previous observations in *Drosophila* S2 cells (6) and indicating evolutionary conservation of DENR functions.

Among the DENR-regulated transcripts were several relevant to cancer, such as *Klhdc8a, Etaa1, Map2k5, Slc20a1* or *Vegfd* (39, 40, 41, 42, 43). Individual inspection confirmed their reduced CDS translation efficiency (Fig. 3A), as well as a shift of ribosome occupancy from the CDS to the 5’ UTR in *Denr*-depleted cells (Fig. 3B). This signature was characteristic of many differential TE genes (see Supplementary Fig. 5A, B), and correlated with the presence of translated uORFs (Supplementary Fig. 5C). Among the strongly DENR-regulated cases, we also noted *Atf5* (Supplementary File 1), whose translation (similar to that of *Atf4*) is known to respond to cellular stress signals via two uORFs, the first of which is reinitiation-permissive whereas the second overlaps the main CDS (35, 44). In Denr-depleted cells, translation of the first uORF (on *Atf5* and, to lesser extent, on *Atf4*) remained unchanged compared to control cells, while translation of the second uORF and the CDS were decreased (Supplementary Fig. 5D, E). These findings validate that our approach indeed delivered reinitiation-prone transcripts.

**Figure 3.**
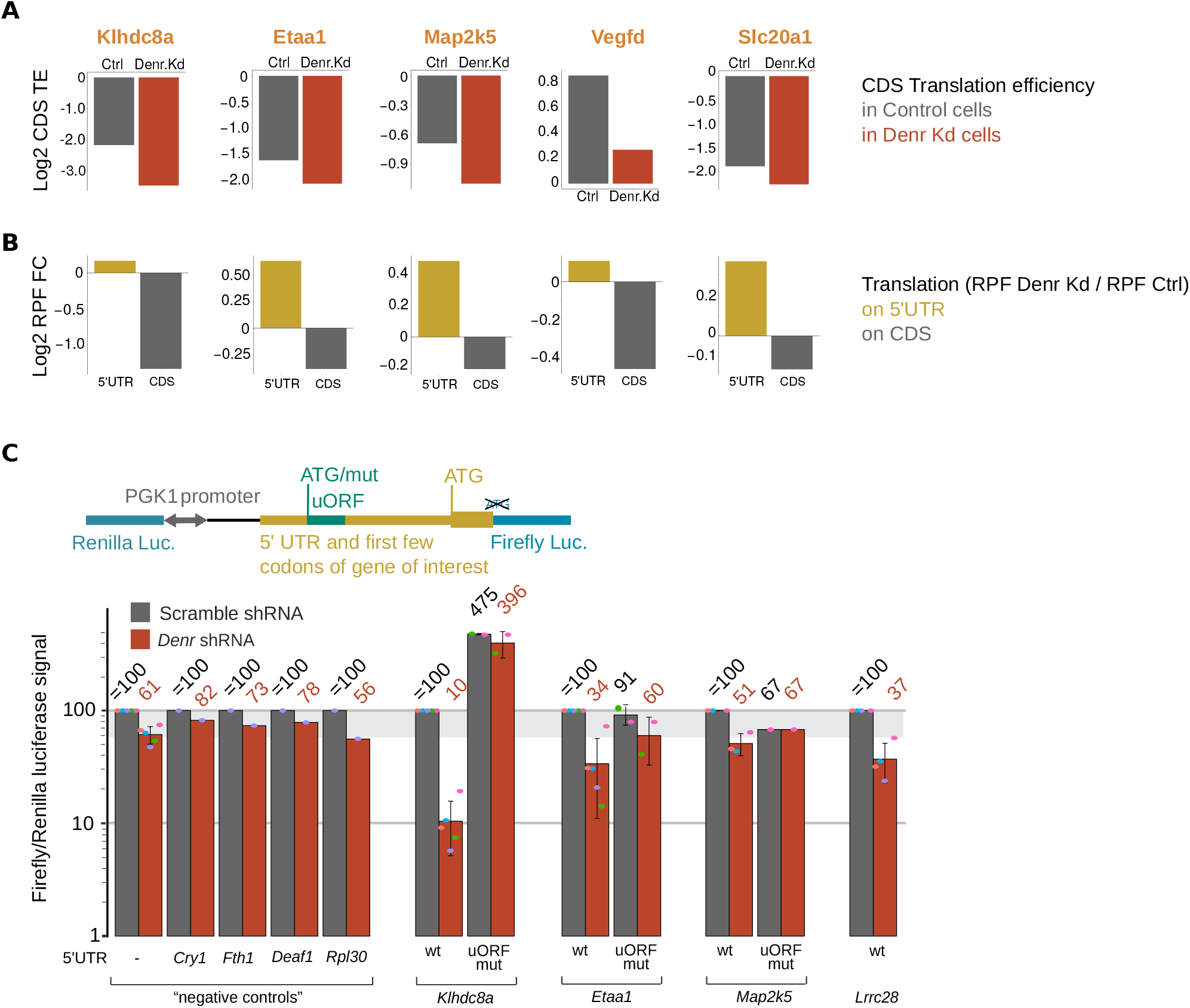
Mutation of uORF start codons relieves DENR dependence. **A)** Representative examples of genes with differential TE. Panels show average CDS translation efficiency in control (grey) and *Denr* Kd (red) cells. **B)** *Denr* Kd-to-control translation on the 5’ UTR (brown) and on the CDS (grey) for the same examples as in (A). DENR targets showed decreased CDS TE (A) and increased 5’ UTR translation concomitant with reduced CDS translation in Denr-deficient cells. **C)** (*Upper*) Schematic representation of the dual luciferase reporter construct, where the 5’ UTR of the gene of interest – with the WT or mutated uORF start – and first few codons of the CDS were cloned upstream of firefly luciferase (FL). (*Lower*) Results of dual luciferase assays, where the influence of the 5’ UTR (WT and uORF mutant) on the expression of FL was measured in control and *Denr* knockdown cells, and internally normalised to RL (expressed from the same bidirectional promoter). Note that for all targets tested, depletion of *Denr* reduced the FL/RL luciferase signal, indicating DENR-dependence of reporter protein expression. Mutation of uORF start codons relieved the inhibition and partially abolished DENR-dependence. For each 5’ UTR assayed, FL/RL signals were normalised to the Scr-transduced non-mutated construct. The area of grey shading marks the non-specific effect of *Denr* loss-of-function in this assay, as judged from the reduction in signal seen for negative control reporters that contained no cloned 5’ UTR (−), or irrelevant 5’ UTRs selected from non-targets.

In order to confirm the DENR- and uORF-dependence of CDS translation for selected presumed targets, we cloned the 5’ UTR and immediate start codon context of several transcripts of interest in frame and upstream of Firefly luciferase (FL) in a lentiviral vector that also expresses *Renilla* luciferase (RL, for internal normalisation) from the same bidirectional promoter (Fig. 3C, upper panel). *Denr* depletion led to a 18-44% reduction in FL/RL signals for five different control reporters that contained either no cloned 5’ UTR, or 5’ UTRs of genes that were not among the differential TE genes (of these, *Cry1* and *Fth1* harbour translated uORFs as well, whereas *Deaf1*, and *Rpl30* do not) (Fig. 3C, lower panel). By contrast, we observed a stronger reduction in FL/RL signals – of 49% to 90% – for four 5’ UTR reporters of presumed DENR target genes (*Etaa1, Klhdc8a, Map2k5* and *Lrrc28*), indicating that DENR was indeed involved in their regulation. Mutating uORF start codons increased reporter signal and abolished most DENR-dependence (Fig. 3C, bottom), indicating that uORF translation was necessary for DENR dependance. Furthermore, in line with the view of DENR as a reinitiation factor, rather than a uORF skipping factor, the presence of a functional stop codon was required for DENR regulation (Supplementary Fig. 5F). uORF-dependent regulation of *Klhdc8a* translation was particularly striking, with a >4-fold translation upregulation in WT cells with a uORF-less construct (Fig. 3C, bottom). We next wished to investigate whether uORF-mediated translational regulation could also be recapitulated *in vivo* and selected *Klhdc8a* for this experiment, as this mRNA contained the strongest, reinitiation-dependent uORF in the cellular assay. We injected mice intravenously with *in vitro*-synthesized mRNAs comprising the *Klhdc8a* 5’ UTR (WT or uORF start codon-mutated) fused to Firefly luciferase CDS and carrying a synthetic polyA tail. We then monitored expression by bioluminescence live imaging in mice. Bioluminescence signal was primarily detected in the livers of the animals, and was 2 – 4.3 times higher in the uORF-less compared to the WT constructs (Supplementary Fig. 6A, B), demonstrating the strong inhibitory activity of the *Klhdc8a* uORF also *in vivo*.

Taken together, we concluded that DENR was involved in the regulation of protein biosynthetic output, and that a set of >200 transcripts containing uORFs was particularly sensitive to loss of this reinitiation factor.

### Linear regression analysis correlates DENR-dependence with uORF strength and with intercistronic distance

uORFs can operate through a bewildering diversity of mechanisms that act on CDS translation (by affecting start site selection/leaky scanning/re-initiation/ribosome shunting), transcript abundance (by eliciting nonsense-mediated mRNA decay), or in some cases through uORF-encoded bioactive peptides (reviewed in (3)). However, rules to predict specific mechanisms from specific uORF features have not yet been established. Our datasets could potentially enable the identification of signatures that render certain uORF-containing transcripts DENR-dependent. Previous studies have suggested that, at least in the context of synthetic reporters in *Drosophila* S2 (6) and human HeLa (7) cells, very short uORFs (coding for 1-2 amino acids) with favourable Kozak contexts were most strongly dependent on DENR for efficient reinitiation at a downstream reporter CDS.

We used multiple regression analysis to determine whether DENR-dependent translation events were associated with specific transcript features. We modeled the CDS TE change observed upon *Denr* knockdown using nine relevant transcript features, namely the number of uORFs, uORF start codon identity (AUG vs. near-cognate) and context, the length, GC content and location of the uORF within the 5’ UTR, the presence of overlapping uORFs, and CDS start codon context (Fig. 4A). The model was applied to all translated uORF-containing genes (N = 4741) and yielded several features with significant predictive value for DENR-sensitivity of TE (Fig. 4B-J; adjusted R^2^ = 0.057, F-statistic = 33.21 on 4731 DF, p < 2.2e-16). Three of the identified features reflected the expected positive relationship between efficient uORF translation and a greater need for reinitiation to subsequently translate the CDS as well. These features were: the number of uORFs present on a transcript (Fig. 4B, b = −0.019, p < 2e-16), the presence of AUG, rather than non-canonical uORF start codons (Fig. 4C, b = −0.05, p < 1.15e-9), and the strength of Kozak contexts around uORF start codons (Fig. 4D, b = −0.0026, p = 0.013). Relatedly, DENR targets were enriched for AUG uORFs as compared to the ensemble of translated uORF-containing transcripts (p = 0.0044, Fisher test, Supplementary Fig. 7A), and genes harboring exclusively AUG uORFs showed the strongest effect on CDS TE in *Denr*-deficient cells (Supplementary Fig. 7B). By contrast, the Kozak context around the CDS start codon bore no predictive value (Fig. 4E, b = −0.0006, p = 0.38), suggesting no DENR involvement in its recognition. Moreover, the presence of CDS-overlapping uORFs was not associated with DENR-dependent TE change (Fig. 4F, b = 0.005, p = 0.44), as expected for a reinitiation factor, given that overlapping uORFs would be reinitiation-incompetent unless the transcripts contained a downstream alternative CDS start codon.

**Figure 4.**
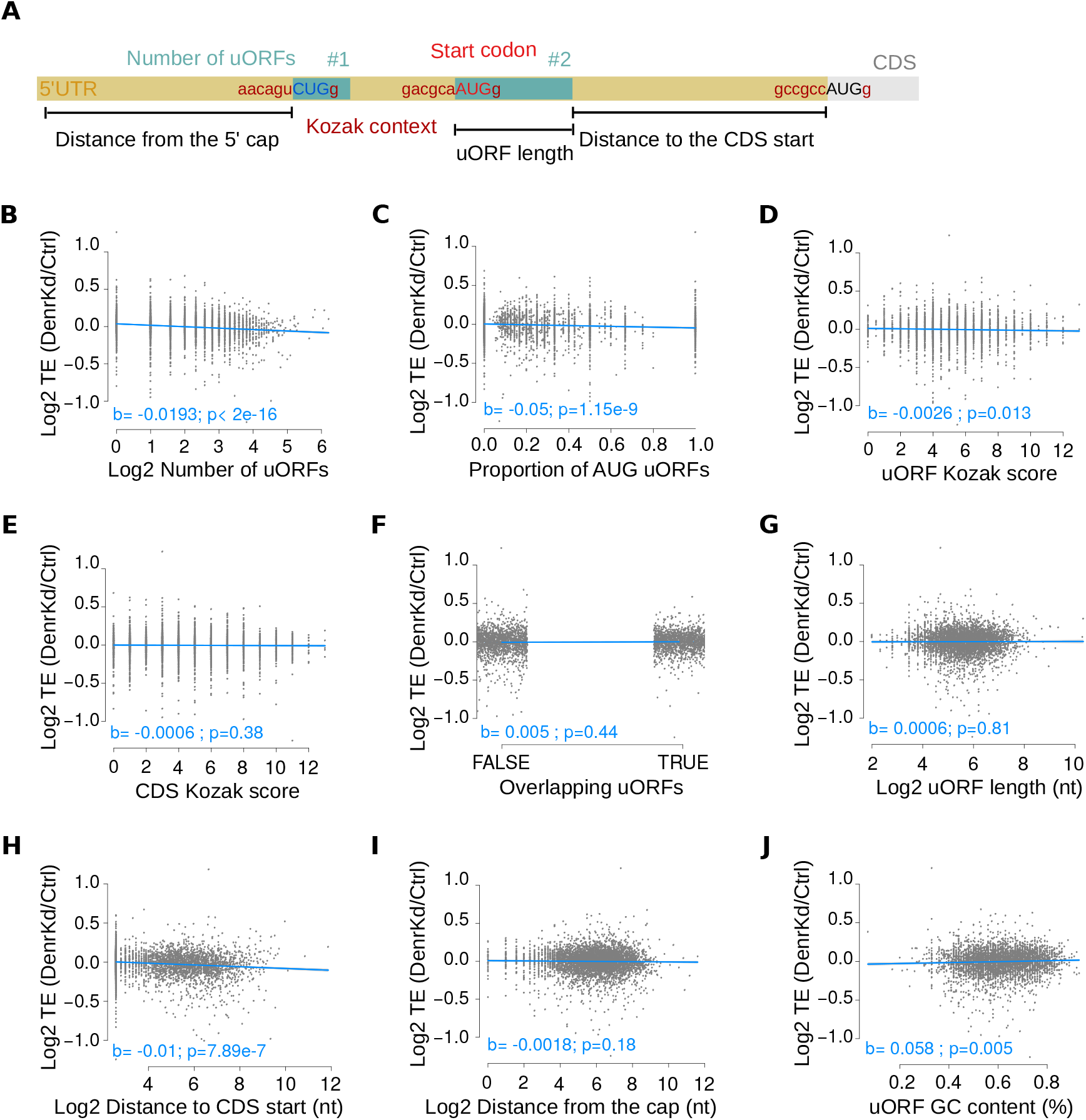
5’ UTR features predictive of translation efficiency regulation by DENR. **A)** Schematic of the 5’ UTR features used as predictors for the modeling of CDS TE change. The number of uORFs, the start codon identity and context, the presence of overlapping uORFs, the uORF length and GC content, and the relative location within the 5’ UTR were considered. Distance to the 5’ cap was computed for the first uORF, and distance to the CDS start was quantified for the last uORF. **B-J)** Partial regression plots for the modeling of translation efficiency fold change (*Denr* Kd vs. Ctrl) on each uORF feature. Indicated in blue are the slope coefficients for the partial regression line and their significance levels (t-test). The number of uORFs (B), the proportion of AUG uORFs (C) and the distance of the last uORF to the CDS start (H) were the strongest predictors.

Two noteworthy outcomes from the model were that uORF length lacked a relationship (Fig. 4G, b = 0.0006, p = 0.81), whereas the uORF-CDS intercistronic distance had predictive value for DENR-sensitivity of CDS TE (Fig. 4H, b = −0.0101, p = 7.89e-7). Briefly, already early studies on a model transcript had suggested that the location of a uORF with respect to the main ORF could dictate the efficiency of reinitiation, as longer spacing would generally allow the reacquisition of initiation factors to the scanning ribosome (45). Our model provides an unbiased and transcriptome-wide validation of this notion for DENR-regulated transcripts, where larger distances from the stop codon of the 3’-most uORF to the start codon of the main ORF were associated with downregulation of TE (Fig. 4H); uORFs located further upstream from the CDS were thus more likely to reinitiate in a DENR-dependent fashion. Importantly, this effect of uORF-CDS distance was not caused by longer 5’ UTRs *per se*, given that the distance from the 5’ cap to the first uORF was not associated with DENR regulation (Fig. 4I, b = −0.0018, p = 0.18). Of note, the uORF-CDS distance was only weakly correlated with 5’ UTR length (Spearman rho = 0.16), whereas the correlation of the 5’ cap-first uORF distance with 5’ UTR length was more prominent (Spearman rho = 0.27, Supplementary Fig. 7C, D).

No linear relationship was found between uORF length and CDS TE (Fig. 4G, b = 0.0006, p = 0.81), in agreement with previous results in human cells using synthetic uORFs encoding >2 amino acids (9 nt) (7). Of note, a length dependence for extremely short uORFs of 1-2 amino acids, as reported by Schleich et al. 2017 (7), would go unnoticed in our analyses, as we excluded them due to the uncertainties associated with their annotation (see Materials and Methods).

Finally, the uORF GC content showed a significant relationship with TE fold change (Fig. 4J, b = 0.058, p = 0.005), suggesting that potential weaker secondary structures (low GC) favoured reinitiation involving DENR.

For the above model, we used the CDS TE fold change as response, which has the benefit of accounting for DENR regulation in a continuous manner. Of note, however, also a logistic regression model based on a binary response (i.e. DENR target as defined in Fig. 2 vs. non-target) recapitulated the main findings (Supplementary Fig. 7E), further supporting our predictions for the strongest cases of DENR translational control. In summary, although the model explained only a relatively modest proportion of the total variance (see Discussion), our results indicated that DENR was important to enable efficient protein synthesis when the transcripts contained uORFs initiating with AUG in strong Kozak contexts (i.e. uORFs expected to be highly translated), and that the distance between the most downstream uORF and the CDS start was an important parameter for efficient reinitiation.

### DENR-mediated reinitiation regulates CLOCK protein biosynthesis

We recently found that uORF translation was pervasive within circadian core clock transcripts, and that the depletion of *Denr* in NIH3T3 cells engendered robust circadian period shortening by 1-1.5 hours (9). We therefore sought to uncover which core clock transcripts were DENR-regulated, and to map the relevant uORFs. Conceivably, these would represent likely contributors to the observed circadian phenotype.

The expression of several core clock genes was altered in *Denr*-depleted vs. control cells. In most cases (*Per1/2/3, Cry1/2, Nr1d1, Rorα*) the observed changes occurred already at the level of mRNA abundance and subsequently led to comparable changes at the footprint level as well (Fig. 5A, and Supplementary Fig. 8A). None of these transcripts (nor *Bmal1*, whose lower RNA abundance was partially compensated at the footprint level) was associated with marked TE reduction (Fig. 5B, and Supplementary Fig. 8B), and we thus considered them unlikely candidates for DENR targets. For *Clock*, however, the reduction in expression was mostly translational, resulting in a ~20% decrease in protein biosynthesis, as quantified from *Clock* RPFs (Figs. 5A,B, and Supplementary Figs. 8A,B). In our initial analysis of differential TE (Fig. 2A), *Clock* had, however, not passed the significance threshold (Supplementary File 1). The reason was that the applied algorithm, Xtail, evaluates statistical significance of TE differences through two separate tests – one regarding differences between the RNA and the RPF fold-change distributions, and a second one evaluating differences in TE distributions directly – and then derives the final p-value from the most conservative of the two tests (22). While the additional test might prevent false positives, it can discard differential TE genes close to the statistical threshold. Importantly, for one of the tests, the TE fold change for *Clock* was significant (p = 0.025 in test 1; p = 0.19 for test 2; Supplementary File 1); *Clock* thus may represent a DENR target whose detection is at the borderline of statistical significance.

**Figure 5.**
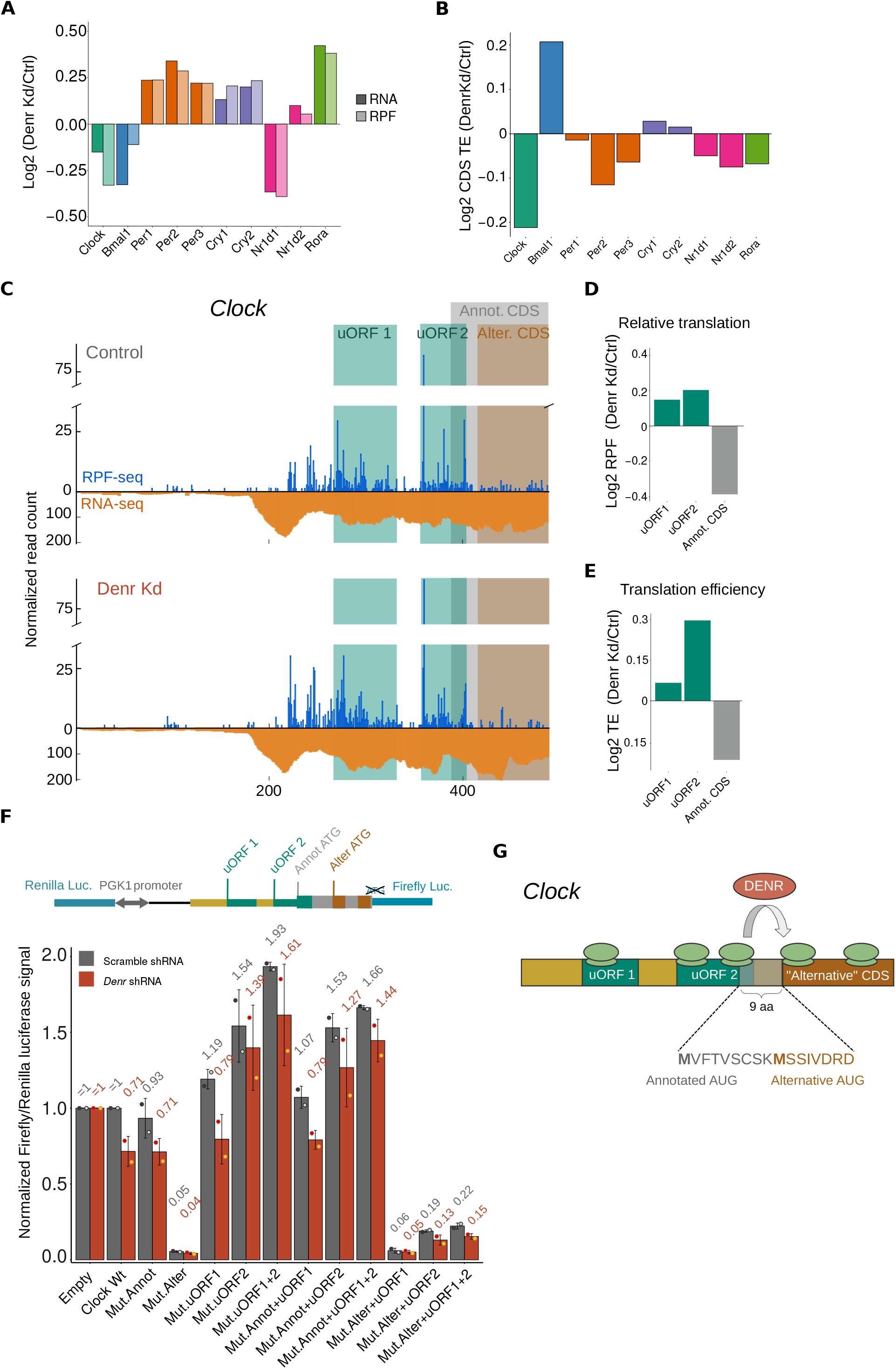
DENR-mediated reinitiation regulates CLOCK protein abundance and N-terminus. **A)** Bar graph of the average RPKM ratio (*Denr* Kd-to-control) for the main circadian core clock genes, at the level of mRNA abundance (dark shadings) and footprint abundance (light shadings). **B)** CDS translation efficiency ratio between *Denr* Kd and control cells showed that *Clock* had a reduced TE upon *Denr* depletion. **C)** Distribution of normalised RPF-seq (blue) and RNA-seq (orange) reads along the 5’ UTR and first 100 nt of *Clock* CDS, in control (top) and *Denr* Kd (bottom) cells. Green shaded areas indicate the two AUG-initiated uORFs; grey area corresponds to the annotated CDS; and brown area to the alternative CDS, in frame with the annotated one. **D)** Quantification of relative translation ratio (*Denr* Kd-to-control) on both uORFs, and on the CDS, showing increased uORF translation and reduced CDS translation in *Denr*-deficient cells. **E)** Quantification of relative translation efficiency (*Denr* Kd-to-control) on both uORFs, and on the CDS showed higher TE on the overlapping uORF (uORF2). **F)** (Top) Schematic representation of the dual luciferase reporter construct, where *Clock* 5’ UTR and first 14 codons of the CDS (containing both the annotated and alternative start codons) were cloned upstream of FL. (Bottom) Dual luciferase assay results for *Clock* WT and mutants. Empty vector contained only the vector-encoded 5’ UTR. FL signal was internally normalised to RL signal; all constructs and conditions were first normalised to the value of condition “empty vector, scr shRNA”; all *Clock* mutant constructs were then additionally normalised to the value from condition “*Clock* WT, scr shRNA”. Signal from all *Denr* shRNA-transduced *Clock* constructs were also normalised to the signal obtained for “empty vector, *Denr*-shRNA”, to remove DENR effects that were not specific to the 5’ UTR. **G)** Schemcatic of how uORFs and reinitiation regulate the translation of the transcript encoding the circadian core clock component *Clock*. *Clock* transcript contains two translated AUG uORFs, the second of which shows high ribosome occupancy and overlaps with the annotated CDS. An additional in-frame AUG is located 9 amino acids downstream of the annotated AUG; usage of the downstream AUG would give rise to a so far unnanotated CLOCK protein variant. Our data suggest that this second AUG is the true start codon for CLOCK biosynthesis in NIH3T3 cells, and that DENR is necessary for translation reinitiation on the alternative CDS after translation of the second uORF.

We annotated two translated AUG-initiated uORFs in *Clock* (Fig. 5C, and Supplementary Fig. 8C-E). The second uORF overlapped with the main CDS in a different frame – a configuration that would render the two translation events (uORF-2 and annotated CDS) mutually exclusive. However, downstream of the uORF-2 stop codon and 10 amino acids into the annotated *Clock* CDS, we noted an additional AUG in the *Clock* coding frame. We thus hypothesised that, when uORF-2 was translated, initiation at this alternative AUG could still allow for the biosynthesis of a CLOCK protein variant that was N-terminally truncated by 9 amino acids (Fig. 5C). In *Denr*-deficient cells, translation was increased on both uORFs (Fig. 5C-D) and, in particular, uORF-2 showed ~23% higher translation rate upon *Denr* depletion (Fig. 5E), suggesting that DENR-mediated reinitiation after uORF-2 was crucial for the translation of the *Clock* CDS.

Using GWIPS-viz, an online genome browser for the visualisation of ribosome profiling data from published studies (46), we looked for further evidence of uORF-2 and alternative CDS translation across mammalian datasets (of note, both annotated and alternative methionine residues are conserved across mammals, see Supplementary Fig. 9). Interestingly, initiating ribosomes can be detected on both uORF-2 and alternative CDS start codons (but not the annotated *Clock* start codon) across human and mouse datasets, and elongating ribosome profiling studies showed good footprint coverage on uORF-2 and the characteristic accumulation of ribosomes at the start and stop codons (Supplementary Fig. 9). Together, the published and our new data thus suggested abundant translation of uORF-2 and of *Clock* CDS from the alternative AUG, indicating that reinitiation has an important role in ensuring CLOCK protein biosynthesis.

In order to investigate uORF usage, effect, and DENR-dependence for CLOCK biosynthesis from the annotated and alternative CDS, we cloned the *Clock* 5’ UTR and first 14 codons (containing both the annotated and alternative AUGs) in frame and upstream of *Firefly* luciferase in our dual luciferase vector (Fig. 5F, upper panel). We then systematically mutated the uORF-1, uORF-2, and CDS annotated/alternative start codons, or combinations thereof (Supplementary Fig. 8F), and compared the effects on luciferase output in *Denr*-depleted vs. control cells. Of note – and in contrast to Fig. 3C – we also adjusted for the effect of *Denr* depletion on the control reporter (whose levels were set to 1) to display the isolated effects of the *Clock* variants.

*Denr* depletion resulted in an almost 30% reduction in expression for the *Clock WT* construct (Fig. 5F, lower panel), in line with the decrease in relative translation quantified for the endogenous transcript (Fig. 5D) and with reduced levels of endogenous CLOCK protein (Supplementary Fig. 10). Surprisingly, mutation of the annotated CDS initiation codon did not substantially affect reporter levels and DENR dependence when compared to *Clock WT*. However, mutation of the alternative AUG decreased luciferase signal by ~95% and abolished DENR dependence (Fig. 5F, lower panel), suggesting that most CLOCK biosynthesis in NIH3T3 cells involves the alternative AUG. Mutation of uORF-1 led to a 19% increase in luciferase signal and fully retained DENR dependence. By contrast, uORF-2 mutation led to a ~50% increase in protein synthesis and eliminated DENR regulation. These findings indicated that uORF-2 translation acts to significantly reduce CLOCK biosynthesis and to render it DENR-dependent. This model was further supported by the combinatorial mutations. Mutation of both uORF-1+2 start codons thus resulted in an additive, >90% increase in protein synthesis, and in a loss of DENR-dependence. In line with minor levels of synthesis of the long CLOCK isoform, mutation of the annotated CDS start codon in combination with uORF mutations led to similar results as mutation of the uORFs alone. Furthermore, mutation of the alternative AUG in combination with uORF mutations led to strongly decreased luciferase signals, consistent with the predominant synthesis of the short CLOCK isoform. In particular, the case where both the alternative CDS and uORF-2 (or uORF-1+2) start codons were mutated – a scenario that would allow unhindered CLOCK synthesis from the annotated AUG – resulted in a >80% reduction in CLOCK protein production (Fig. 5F, lower panel). Taken together, we concluded that *Clock* required DENR for efficient protein synthesis from a CDS that is 9 codons shorter than the annotated one, and that uORF-2 plays a key role in coordinating the complex interactions that occur at the translational level on the *Clock* transcript (Fig. 5G). Moreover, we propose that the CLOCK N-terminus is likely wrongly annotated not only in NIH3T3 cells, but based on our analysis of published ribosome profiling data, also in other cell types and in tissues (Supplementary Fig. 9).

## DISCUSSION

The annotation of translated uORFs has remained a particular challenge, because they are short, frequently initiate from non-AUG codons, and do not obey the same rules of evolutionary conservation as most protein coding sequences (3). This has changed with the availability of ribosome profiling data, which has made it possible to systematically identify translated uORFs from footprints. The resulting burst in interest in uORFs and the rising appreciation of how prevalent their usage is in mammals, is, however, not yet matched by an understanding at the mechanistic level. In this study, we have investigated one particularly intriguing mechanism that is implicated in the regulation through uORFs, namely translation reinitiation. As an entry point, we have used the loss-of-function of DENR, a non-canonical initiation factor recently associated with reinitiation both in *Drosophila* (6) and in HeLa cells (7), with the aim of identifying rules that explain why certain uORFs trigger DENR-dependent reinitiation while others do not. The conclusions we were able to draw from experiments in mouse NIH3T3 fibroblasts showed similarities and striking differences to the previous studies in fly and in human cells (6, 7). Species-specificity in uORF biology may explain some of the disparities, but differences in experimental and computational approaches likely have an important influence as well.

We view our quantitative modeling approach of DENR-dependent changes in CDS TE as particularly unbiased as it relies on translational changes detected on endogenous mRNAs transcriptome-wide. Several features carry predictive value for the magnitude of DENR-mediated CDS TE regulation. Of these, three (number of uORFs, start codon identity, start codon sequence context) likely reflect a straightforward logic between a stronger requirement for reinitiation when the usage of uORFs is efficient. This outcome is expected and fully consistent with human (7) and fly data (6) based on reporter assays. Two other outcomes from the model are noteworthy. First, uORFs located further away from the main ORF are more prone to trigger DENR-dependent reinitiation. Such intercistronic distance effects were reported early on in the uORF field, typically using distinct uORF-containing model transcripts (45, 47, 48). They had generally been assumed to reflect the kinetics of association and dissociation of initiation and termination factors. Our data validate this idea in a transcriptome-wide fashion and are in line with the hypothesis that longer distances allow for extended re-scanning times, which would be favourable for the loss of termination factors and/or the re-acquisition of the initiation machinery, on time for CDS translation. Second, it is noteworthy that uORF length lacks predictive value in our model. The hypothesis that increased uORF length impairs reinitiation efficiency hence does not appear valid, at least for DENR-regulated reinitiation events. As a caveat, the nature of our approach (transcriptome-wide from endogenous transcripts) did not allow to confidently annotate uORFs down to only 1-2 amino acids in length. We may thus be missing DENR effects that have recently been observed in reporters carrying such ultra-short synthetic uORFs (7). We can evidently also not exclude that the time required for uORF translation in determining scanning resumption is relevant nevertheless, as linear length *per se* may simply not be a good proxy for translation time (2, 8, 49).

Overall, our model explains a relatively modest proportion of the total variance (adjusted R^2^ of 0.057). This is reminiscent of analogous models that have been developed to predict, for example, miRNA target sites, whose adjusted R^2^s range from 5-13%, but that have nevertheless proven biologically useful (50). In our case, the presence of multiple uORFs on a transcript, often overlapping each other, and each with its own, distinct features (start codon, length, distance to the CDS, etc), and the resulting combinatorial effects, clearly complicated their scoring and likely masked some of the origins of translational regulation. We deliberately wished to keep our model simple, aiming at identifying the most significant predictors, but at the expense of overall explanatory power. In the future, the use of other regression types and models that include interaction terms could potentially explain higher biological variability, although at the cost of becoming considerably more difficult to interpret. Overall, our study constitutes a first attempt to analyse signatures associated with reinitiation through DENR. It was already able to recapitulate previous observations (6, 7), and as we continue to identify more molecular features that can regulate non-canonical translation initiation and that can be integrated into the model in the future, a larger proportion of biological variance will be explained.

We have identified 240 transcripts as particularly DENR-sensitive. Although we lack formal proof that they are direct DENR targets, the observations that (1) most of them show TE downregulation and few upregulation in the absence of DENR, that (2) they are depleted for CDS-overlapping and enriched for non-overlapping uORFs, and that (3) a DENR-dependent inhibitory effect of their 5’ UTRs can be rescued by uORF start codon mutations in reporter assays, all make a direct activity of DENR on these transcripts very likely. Moreover, these observations are all compatible with a role of DENR in reinitiation, but not in uORF skipping (which could have been an alternative interpretation of the redistribution of ribosomes from CDS to 5’ UTRs seen in Fig. 1E). In particular, we would not expect CDS-overlapping uORFs to be depleted (Supplementary Fig. 4B) if DENR acted to regulate targets via uORF skipping. Moreover, mutating the uORF stop codon should not have had an influence on the DENR-effect in our reporter assays, if the activity of DENR consisted in regulating the efficiency of uORF start codon selection (Supplementary Fig. 5F).

Many identified DENR targets are involved in proliferation and associated with cancer. Of note, DENR-MCTS1 have themselves been identified as oncogenes (51) that are upregulated, for example, in prostate cancer (52). Moreover, translational reprogramming, especially with regard to upstream initiation sites, appears to represent an important cancer driver (30), and understanding how the translation of cancer-relevant DENR targets is regulated may thus lead to important insights of diagnostic or therapeutic value. We found, for example, very strong DENR requirement for the translation of Kelch Domain Containing 8A (KLHDC8A). It has been reported that KLHDC8A overexpression in human gliomas that become resistant to epidermal growth factor receptor (EGFR) silencing (a commonly used anticancer therapeutic approach), allows tumours to maintain aggressiveness through an unknown EGFR-independent pathway (39). It is conceivable that upon loss of EGFR sensitivity, KLHDC8A levels increase through DENR-mediated translational control. Similarly, other candidates from our DENR target list (Supplementary file 1) are upregulated in (and possibly causally contributing to) different cancers, such as Vascular endothelial growth factor D (VEGF-D), or Mitogen-Activated Protein Kinase 5 (MAP2K5) (41, 43). By understanding how DENR regulates its targets, it may thus be possible to reduce reinitiation activity and thus the oncogenic levels of the modulated proteins in proliferating, but not in quiescent cells.

A comparison of DENR targets found in our and in previous studies returned little overlap. Of the eight targets validated in HeLa cells (7), only one (*Tmem60*) was expressed in NIH3T3 cells. *Tmem60* was predicted to undergo a 25% translational downregulation upon *Denr* loss-of-function and indeed showed a decrease in that range in reporter assays in HeLa cells (7), whereas in our system a ~8% decrease in CDS TE of endogenous *Tmem60* was quantified. Moreover, for the ensemble of 104 predicted targets in human cells (7), >90% were not expressed in NIH3T3 cells or lacked a mouse orthologue; others (e.g. *Lca5, Mllt11* or *Dsel*) were indeed downregulated upon *Denr* knockdown, though to a lesser extent. Conceivably, DENR targets are species- and cell type-specific; moreover, it is likely that the differences in the magnitude of translational regulation observed across studies are the result of different experimental and analytical approaches.

How do DENR and circadian rhythms mechanistically connect? The DENR target list did not reveal core clock genes that would explain the circadian clock phenotype, i.e. the previously reported ~ 1.5-hour period shortening of free-running oscillations (9). Individual inspection of core clock gene transcripts, however, suggested that *Clock* was a DENR target just below the significance threshold. Follow-up experiments confirmed *Clock* regulation through DENR and allowed mapping the responsible uORF. These findings establish uORF translation and re-initiation as a novel, potentially regulated mechanism acting on core clock gene expression, which opens several intriguing questions. First, is it possible that reduced biosynthesis of *Clock* would be responsible for the observed phenotype? Loss-of-function of a general gene expression machinery component typically goes along with sizeable distortions in global RNA and protein profiles, and *Denr* knockdown is no exception in this respect. Any phenotype thus likely represents a composite effect, rather than the result of a single misregulated gene expression event. We therefore do not view reduced CLOCK biosynthesis as a satisfactory hypothesis that alone would be able to explain the substantial period shortening. Nevertheless, it is reassuring that *Clock^−/−^* fibroblasts have recently been shown to have shorter periods as well (53), which suggests that downregulation of *Clock* translation upon *Denr* depletion may be a factor contributing to the phenotype.

A second intriguing question relates to the alternative initiation site that would generate an N-terminally truncated variant of CLOCK. We propose that the 9 amino acid shorter CLOCK is the main variant produced in NIH3T3 cells, and likely in other cell types and species as well. What are the functional implications? The bHLH domain responsible for DNA binding of CLOCK starts at amino acid 26, and the very N-terminus has not yet been associated with any function of the protein and is also not included in the crystal structure of the CLOCK-BMAL1 complex (54). However, it is noteworthy that for several myogenic bHLH transcription factors, N-terminal motifs are critical for cooperative DNA binding with cofactors (55). Much of our current knowledge on CLOCK stems from *in vivo* and *in vitro* experiments that have used *Clock* cDNA constructs devoid of the endogenous uORF regulation, and thus defining CLOCK’s N-terminus to the annotated one. It will be exciting to re-evaluate selected DNA-binding and activity assays using both CLOCK isoforms side by side.

Finally, a speculative question to ask is whether DENR-mediated control is a regulated, rather than constitutive step in CLOCK biosynthesis, serving for example to uncouple CLOCK protein abundance from mRNA levels in specific cell types or under certain physiological conditions. Of note, there is precedence from *Neurospora crassa* for uORF translation that mediates temperature-dependent regulation of biosynthesis of the core clock component FRQ (56). In the case of DENR, it is interesting that this protein was originally identified because its expression increased in cultured cells at high density, but not during growth arrest *per se*, which led to its name Density Regulated Protein (57). First links between cell density and circadian rhythmicity have indeed been described in recent years (58, 59), and it would be interesting to assess whether CLOCK uORFs and DENR play roles in determining density-dependent differences of clock parameters.

## DATA AVAILABILITY

High-throughput sequencing data was deposited to GEO, under accession number: GSE124793 (SuperSeries composed of ribosome profiling data: GSE116221 and iCLIP data: GSE124790). Code for ribosome profiling pre-processsing, aligning and quantification, and for uORF annotation, is available online at http://doi.org/10.5281/zenodo.521199.

## SUPPLEMENTARY DATA

Supplementary data is available.

## ACKNOWLEDGEMENTS

We thank the Lausanne Genomics Technologies Facility for high-throughput sequencing infrastructure. We thank Angelica Liechti and Georgia Katsioudi who were involved in initial steps of reporter plasmid clonings. We are grateful to Igor Ruiz de los Mozos for assistance with the iClip analyses through iCount.

