## Supplementary material for "Charting DENR-dependent translation reinitiation uncovers predictive uORF features and links to circadian timekeeping via Clock"

A

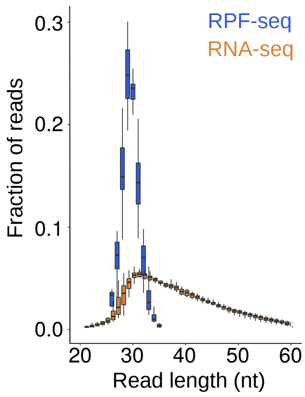


B

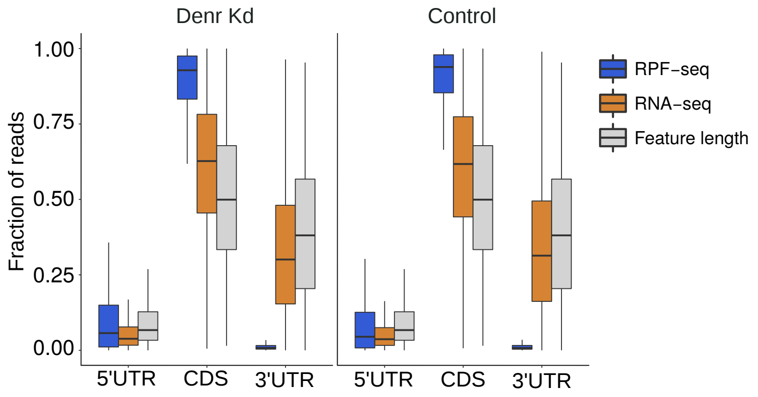


C

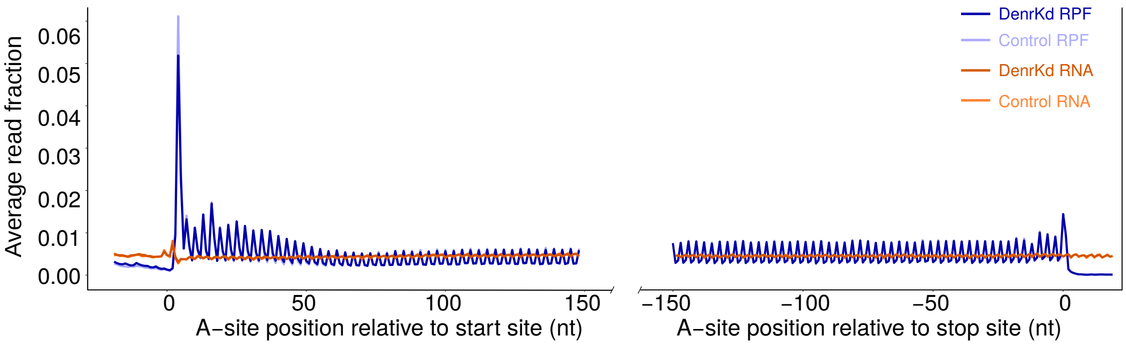


D

Correlation between samples RPF-seq

| | Denr1_1 | Denr1_2 | Denr1_3 | Denr2_1 | Denr2_2 | Denr2_3 | Gfpsh_1 | Gfpsh_2 | Scrsh_1 | Scrsh_2 | Scrsh_3 |
|---|---|---|---|---|---|---|---|---|---|---|---|
| Denr1_1 | 1 | 1 | 0.99 | 1 | 0.99 | 0.99 | 0.99 | 0.99 | 0.99 | 0.98 | 0.97 |
| Denr1_2 | | 1 | 0.99 | 1 | 1 | 0.99 | 0.99 | 1 | 0.99 | 0.99 | 0.98 |
| Denr1_3 | | | 1 | 0.99 | 0.99 | 1 | 0.98 | 0.99 | 0.99 | 0.99 | 0.99 |
| Denr2_1 | | | | 1 | 1 | 0.99 | 0.99 | 1 | 0.99 | 0.99 | 0.98 |
| Denr2_2 | | | | | 1 | 1 | 0.99 | 1 | 0.99 | 0.99 | 0.99 |
| Denr2_3 | | | | | | 1 | 0.99 | 0.99 | 0.99 | 0.99 | 0.99 |
| Gfpsh_1 | | | | | | | 1 | 1 | 0.99 | 0.99 | 0.98 |
| Gfpsh_2 | | | | | | | | 1 | 0.99 | 0.99 | 0.98 |
| Scrsh_1 | | | | | | | | | 1 | 0.99 | 0.98 |
| Scrsh_2 | | | | | | | | | | 1 | 1 |
| Scrsh_3 | | | | | | | | | | | 1 |

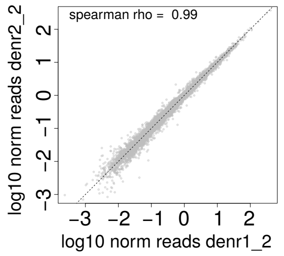


Correlation between samples RNA-seq

| | Denr1_1 | Denr1_2 | Denr1_3 | Denr2_1 | Denr2_2 | Denr2_3 | Gfpsh_1 | Gfpsh_2 | Scrsh_1 | Scrsh_2 | Scrsh_3 |
|---|---|---|---|---|---|---|---|---|---|---|---|
| Denr1_1 | 1 | 0.99 | 0.97 | 0.99 | 0.98 | 0.98 | 0.99 | 0.98 | 0.98 | 0.97 | 0.95 |
| Denr1_2 | | 1 | 0.99 | 0.99 | 1 | 0.99 | 0.99 | 1 | 0.99 | 0.99 | 0.97 |
| Denr1_3 | | | 1 | 0.99 | 0.99 | 0.99 | 0.98 | 0.99 | 0.99 | 0.99 | 0.99 |
| Denr2_1 | | | | 1 | 0.99 | 0.99 | 1 | 0.99 | 0.99 | 0.99 | 0.97 |
| Denr2_2 | | | | | 1 | 1 | 0.99 | 1 | 0.99 | 1 | 0.98 |
| Denr2_3 | | | | | | 1 | 0.99 | 0.99 | 0.99 | 0.99 | 0.98 |
| Gfpsh_1 | | | | | | | 1 | 1 | 0.99 | 0.99 | 0.97 |
| Gfpsh_2 | | | | | | | | 1 | 0.99 | 0.99 | 0.98 |
| Scrsh_1 | | | | | | | | | 1 | 1 | 0.98 |
| Scrsh_2 | | | | | | | | | | 1 | 0.98 |
| Scrsh_3 | | | | | | | | | | | 1 |

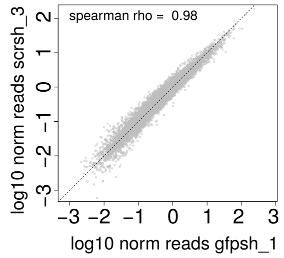


E

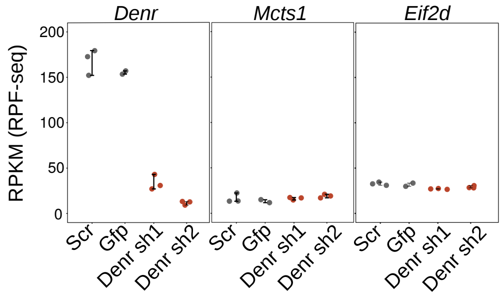


F

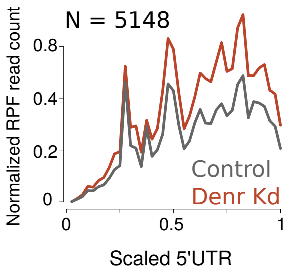


G

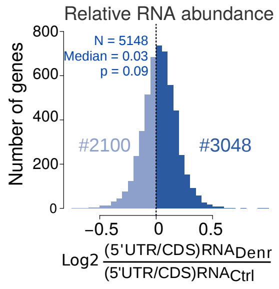

**Supplementary Figure 1. Ribosome profiling in Denr knockdown NIH3T3 mouse fibroblasts.** **A)** Read length distribution for ribosome footprint (RPF-seq) and mRNA (RNA-seq). Footprint length distribution peaks sharply at 29-30 nt, whereas RNA-seq reads showed a broader distribution, as expected. **B)** Read mapping to transcript features. RPF-seq and RNA-seq compared to a distribution expected from the relative feature size (grey). In both *Denr* Kd and control cells, RPF-seq but not RNA-seq reads were depleted from 3′ UTRs and enriched on the CDS. A significant amount of reads also mapped to 5′ UTRs. **C)** Metagene analysis of the footprints' ribosome A-site positions relative to the start and stop codons of the transcripts. Read density at each position was averaged across genes with a single protein-coding isoform, with a CDS RPF RPKM $>$ 5 and longer than 400 nt ($n = 3141$). This analysis revealed the trinucleotide periodicity of RPF-seq, but not RNA-seq, reads. **D)** Tables of Spearman correlation coefficients between samples for normalized CDS reads. On the right, one representative example of normalised CDS reads correlation between replicate samples. **E)** RPKM values for CDS-mapping RPF-seq read counts for *Denr* (left), *Mcts1* (middle), and *Eif2d* (right). Each dot corresponds to a sample and replicate and error bars indicate mean$\pm$sd. **F)** Metagene analysis of normalized RPF-seq read counts across 5′ UTRs. 5′ UTRs were scaled to forty windows of equal size and RPF signal was averaged within each window and across genes. *Denr*-deficient cells showed an accumulation of ribosome footprints on the 5′ UTR – possibly due to the presence of translated uORFs – that can explain the change in relative 5′ UTR shown in Fig. 1E, and indicative of a translational landscape remodelling. **G)** Histogram of 5′ UTR-to-CDS mRNA abundance ratios (from RNA-seq) between *Denr* Kd and control cells, using same genes as in Fig. 1E. In contrast to translation, relative 5′ UTR transcript levels did not change between conditions (median = 0.03, $p = 0.09$).

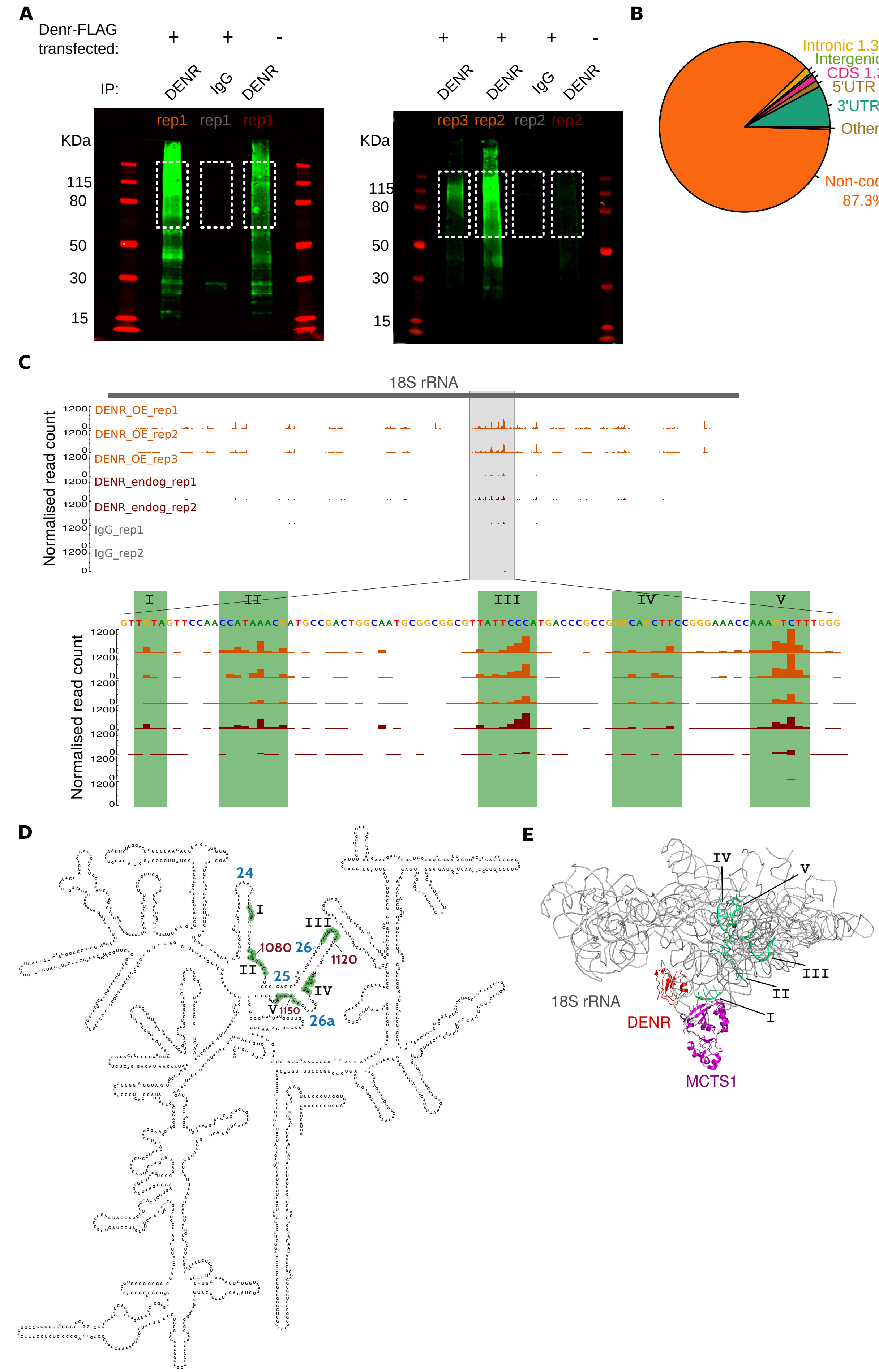
A
Denr-FLAG transfected:
+ + -
+ + + -
IP:
DENR IgG DENR
DENR DENR IgG DENR
rep1 rep1 rep1
rep3 rep2 rep2 rep2
KDa
115
80
50
30
15
B
Intronic 1.3%
Intergenic 0.4%
CDS 1.3%
5'UTR 1.3%
3'UTR 7.9%
Other 0.5%
Non-coding 87.3%
C
18S rRNA
Normalised read count
DENR_OE_rep1
DENR_OE_rep2
DENR_OE_rep3
DENR_endog_rep1
DENR_endog_rep2
IgG_rep1
IgG_rep2
1200
0
I
II
III
IV
V
GTTGTAGTTCCAACCATAAACGATGCCGACTGGCAATGCGGCGGCGTTATTCCCATGACCCGCCGGGCAGCTTCCGGGAAACCAAAGTCTTTGGG
D
24
25
26
26a
1080
1120
1150
E
18S rRNA
DENR
MCTS1

**Supplementary Figure 2. DENR-MCTS1 iCLIP in NIH3T3 cells. A)** Li-Cor image of infrared adaptor-ligated protein-RNA complexes. The region within the white rectangle – likely representing RNA-bound DENR:MCTS1 (together ∼50KDa) complexes – was excised and used for library preparation. NIH3T3 cells overexpressing DENR were used in triplicate for iCLIP experiments; non-overexpressing cells in duplicate, and an IgG IP was done in duplicate, as a negative control. **B)** Mapping summary of iCLIP experiments in cells overexpressing DENR. Sections represent percentage of cDNA read counts per kilobase uniquely mapping to the regions indicated. **C)** Overview of high-confidence binding peaks mapping within the 18S rRNA (top), and zoom into the region with the five most prominent peaks (bottom), located within helixes 24-26. Plots were generated with Biodalliance genome visualisation tool (biodalliance.org). **D)** Secondary structure of 18S rRNA; in green, the five most prominent binding sites highlighted in C. **E)** Crystal structure of DENR:MCTS1 in complex with 18S rRNA (PDB: 5VYC; from Lomakin et al. 2017, ref. 32), highlighting the five most prominent binding sites, as in C,D.

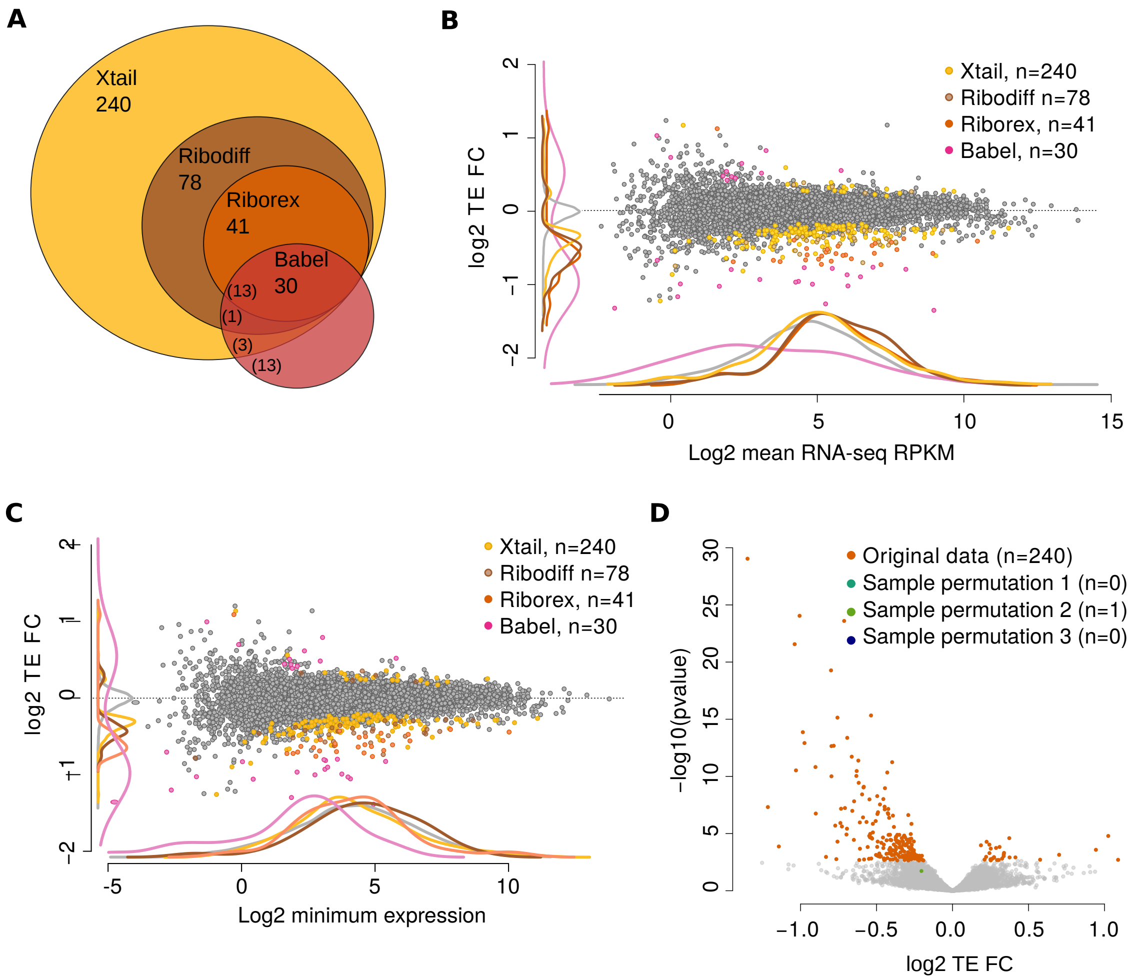
A
Xtail
240
Ribodiff
78
Riborex
41
Babel
30
(13)
(1)
(3)
(13)
B
Xtail, n=240
Ribodiff n=78
Riborex, n=41
Babel, n=30
log2 TE FC
Log2 mean RNA-seq RPKM
C
Xtail, n=240
Ribodiff n=78
Riborex, n=41
Babel, n=30
log2 TE FC
Log2 minimum expression
D
Original data (n=240)
Sample permutation 1 (n=0)
Sample permutation 2 (n=1)
Sample permutation 3 (n=0)
−log10(pvalue)
log2 TE FC

**Supplementary Figure 3. Comparison of algorithms for differential translation efficiency analysis. A)** Venn diagram of differential TE genes detected by Xtail (n=240), RiboDiff (n=78), Riborex (n=41) or Babel (n=30) with an FDR $<$ 0.1. The analysis showed agreement among the first three algorithms, with Xtail detecting the largest number of significant cases. **B)** Scatter plot with marginal densities of average mRNA abundance vs. TE fold change (*Denr* Kd/Ctrl), with differential TE genes highlighted for each algorithm. For cases detected by several algorithms, the genes are coloured according to the method detecting fewer genes, and density lines correspond to the distribution of all detected cases. The majority of Xtail, RiboDiff and Riborex detected genes showed decreased TE in *Denr* deficient cells as expected, and spanned all levels of expression, whereas Babel detected similar numbers of up- and downregulated TE genes and showed a bias for lowly expressed genes. **C)** As B, but showing minimal expression in the x-axis, i.e. the lowest value among all RNA-seq and RPF-seq replicates. **D)** Volcano plot of differential TE genes detected by Xtail on the original data (n=240) and on three condition-shuffled datasets. Sample permutation yielded nearly no significant genes, indicating low false positive detection by Xtail.

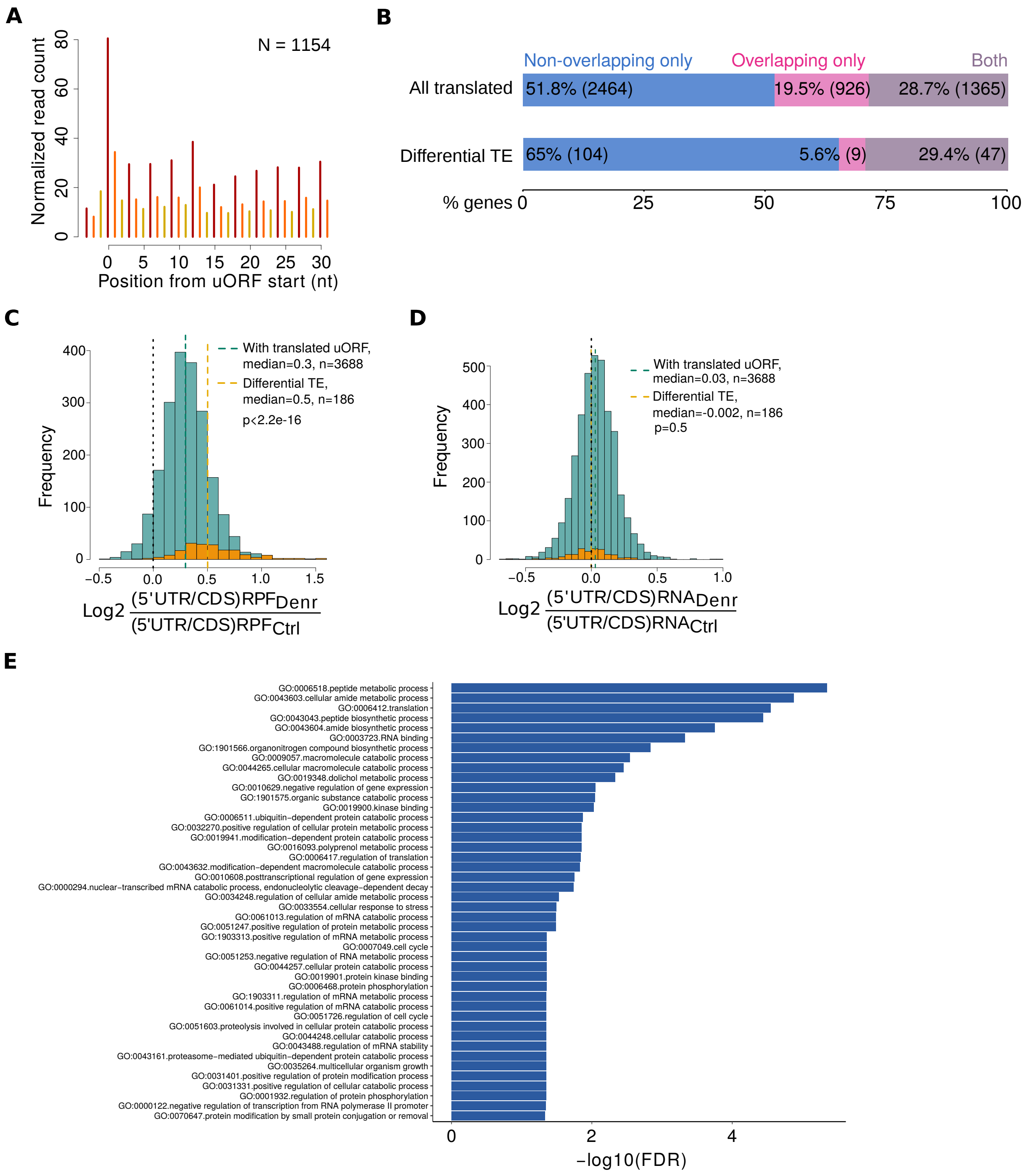

A
N = 1154
Normalized read count
Position from uORF start (nt)
B
Non-overlapping only
Overlapping only
Both
All translated
51.8% (2464)
19.5% (926)
28.7% (1365)
Differential TE
65% (104)
5.6% (9)
29.4% (47)
% genes
C
With translated uORF, median=0.3, n=3688
Differential TE, median=0.5, n=186
p<2.2e-16
Frequency
Log2 (5'UTR/CDS)RPF Denr / (5'UTR/CDS)RPF Ctrl
D
With translated uORF, median=0.03, n=3688
Differential TE, median=-0.002, n=186
p=0.5
Frequency
Log2 (5'UTR/CDS)RNA Denr / (5'UTR/CDS)RNA Ctrl
E
GO:0006518.peptide metabolic process
GO:0043603.cellular amide metabolic process
GO:0006412.translation
GO:0043043.peptide biosynthetic process
GO:0043604.amide biosynthetic process
GO:0003723.RNA binding
GO:1901566.organonitrogen compound biosynthetic process
GO:0009057.macromolecule catabolic process
GO:0044265.cellular macromolecule catabolic process
GO:0019348.dolichol metabolic process
GO:0010629.negative regulation of gene expression
GO:1901575.organic substance catabolic process
GO:0019900.kinase binding
GO:0006511.ubiquitin-dependent protein catabolic process
GO:0032270.positive regulation of cellular protein metabolic process
GO:0019941.modification-dependent protein catabolic process
GO:0016093.polyprenol metabolic process
GO:0006417.regulation of translation
GO:0043632.modification-dependent macromolecule catabolic process
GO:0010608.posttranscriptional regulation of gene expression
GO:0000294.nuclear-transcribed mRNA catabolic process, endonucleolytic cleavage-dependent decay
GO:0034248.regulation of cellular amide metabolic process
GO:0033554.cellular response to stress
GO:0061013.regulation of mRNA catabolic process
GO:0051247.positive regulation of protein metabolic process
GO:1903313.positive regulation of mRNA metabolic process
GO:0007049.cell cycle
GO:0051253.negative regulation of RNA metabolic process
GO:0044257.cellular protein catabolic process
GO:0019901.protein kinase binding
GO:0006468.protein phosphorylation
GO:1903311.regulation of mRNA metabolic process
GO:0061014.positive regulation of mRNA catabolic process
GO:0051726.regulation of cell cycle
GO:0051603.proteolysis involved in cellular protein catabolic process
GO:0044248.cellular catabolic process
GO:0043488.regulation of mRNA stability
GO:0043161.proteasome-mediated ubiquitin-dependent protein catabolic process
GO:0035264.multicellular organism growth
GO:0031401.positive regulation of protein modification process
GO:0031331.positive regulation of cellular catabolic process
GO:0001932.regulation of protein phosphorylation
GO:0000122.negative regulation of transcription from RNA polymerase II promoter
GO:0070647.protein modification by small protein conjugation or removal
-log10(FDR)

**Supplementary Figure 4. Analysis of differential TE genes. A)** Triplet periodicity of ribosome footprints within translated uORFs. Genes containing a single uORF were included in the analysis (n = 1154 genes/uORFs) to avoid uORFs overlapping to each other. **B)** Proportion of translated uORF-containing genes – specified as non-overlapping uORFs only, overlapping uORFs only, or both – transcriptome-wide (all translated) and among DENR targets (differential TE). The latter showed a depletion of overlapping uORFs and enrichment for genes containing only non-overlapping uORFs. **C)** Histogram of 5′ UTR-to-CDS translation between *Denr* Kd and control cells, for all uORF-containing genes (grey, n=3688) and for differential TE genes (orange, n=186). Genes with a minimum of 20 RPF-seq reads in the 5′ UTR in 3/4 of samples were included in the analysis. **D)** Same as in (C) for RNA-seq reads. Differential TE genes showed increased relative 5′ UTR translation, but equal transcript levels, in *Denr*-deficient cells, that was higher than for the whole set of uORF-containing genes, suggestive of impaired translation reinitiation among targets. **E)** Gene Ontology (GO) analysis for Biological process, Metabolic function and Reactome functional categories of genes with differential TE. Bars show -log10 for categories with an FDR $< 0.05$.

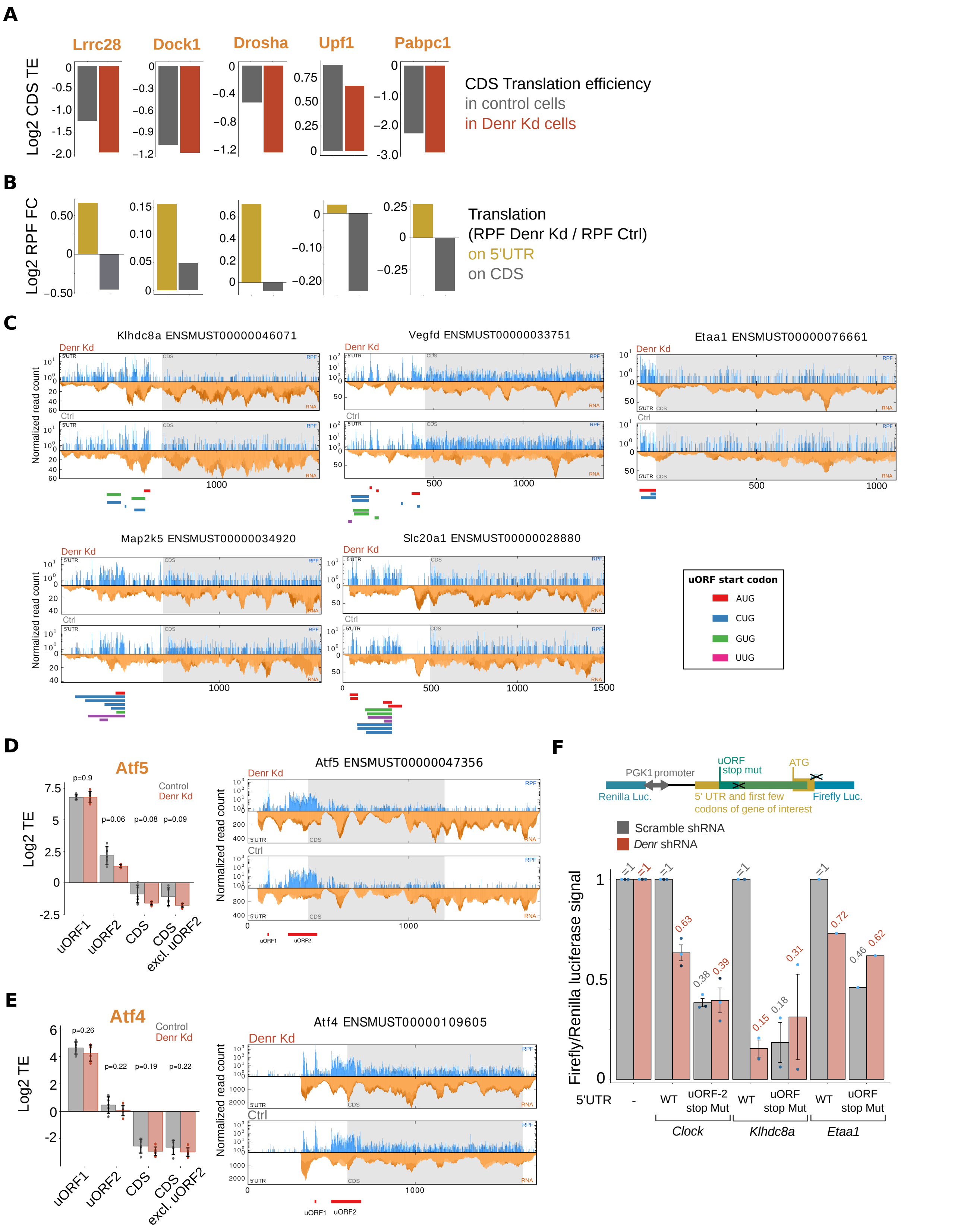

A
Lrrc28
Dock1
Drosha
Upf1
Pabpc1
Log2 CDS TE
CDS Translation efficiency
in control cells
in Denr Kd cells
B
Log2 RPF FC
Translation
(RPF Denr Kd / RPF Ctrl)
on 5'UTR
on CDS
C
Klhdc8a ENSMUST00000046071
Vegfd ENSMUST00000033751
Etaa1 ENSMUST00000076661
Map2k5 ENSMUST00000034920
Slc20a1 ENSMUST00000028880
Denr Kd
Ctrl
Normalized read count
uORF start codon
AUG
CUG
GUG
UUG
D
Atf5
Log2 TE
Control
Denr Kd
uORF1
uORF2
CDS
CDS excl. uORF2
Atf5 ENSMUST00000047356
E
Atf4
Atf4 ENSMUST00000109605
F
PGK1promoter
uORF stop mut
ATG
Renilla Luc.
5' UTR and first few codons of gene of interest
Firefly Luc.
Scramble shRNA
Denr shRNA
Firefly/Renilla luciferase signal
5'UTR
WT
uORF-2 stop Mut
uORF stop Mut
Clock
Klhdc8a
Etaa1

**Supplementary Figure 5. Examples of differential TE genes. A)** Representative examples of genes with differential TE. Panels show average CDS translation efficiency in control (grey) and *Denr* Kd (red) cells. **B)** *Denr* Kd-to-control translation on the 5′ UTR (brown) and on the CDS (grey). DENR targets showed decreased CDS TE (A) and increased 5′ UTR translation concomitant with reduced CDS translation in *Denr*-deficient cells. **C)** Normalized RPF (blue) and RNA (orange) read count distribution on several DENR targets, in knockdown (top) and control (bottom) cells. Different blue and orange shadings correspond to each replicate per condition. The first 1000 nt of the CDS are shown. Coloured boxes below profiles indicate translated uORFs. **D)** (Left) Quantification of translation efficiency on *Atf5* uORFs and main CDS, in control (grey) and *Denr*-deficient (red) cells. Dots represent replicate samples, error bars indicate mean $\pm$ sd, and p-values are derived from t-test. (Right) As in (C), for *Atf5*. **E)** As in (D), for *Atf4*. **F)** (Top) Schematic representation of the dual luciferase reporter construct, in which the uORF stop codon was mutated so that the uORF translates into the CDS, in a different frame. These constructs allow testing the dependence on a functional stop codon for DENR-mediated regulation. Of note, if the function of DENR consisted in regulating the efficiency of uORF start codon selection, mutating the uORF stop codon should not have an influence on the DENR-effect in the reporter assay. (Bottom) Results of dual luciferase assay in control (grey) and *Denr*-depleted cells (red). Removal of the stop codons of *Clock*, *Klhdc8a* and *Etaa1* uORFs led to loss of DENR-dependence, indicating that the presence of a stop codon is necessary for DENR action, in line with a role of DENR as a reinitiation factor rather than a leaky scanning/uORF skipping factor. FL signal was internally normalised to RL expressed from the same bidirectional promoter. FL/RL signals were normalised to the Scr-transduced non-mutated ("WT") vector. Signals from all *Denr*-shRNA transduced constructs were also normalised to the signal obtained for "empty vector, *Denr*-shRNA", to remove DENR effects that were not specific to the 5′ UTR.

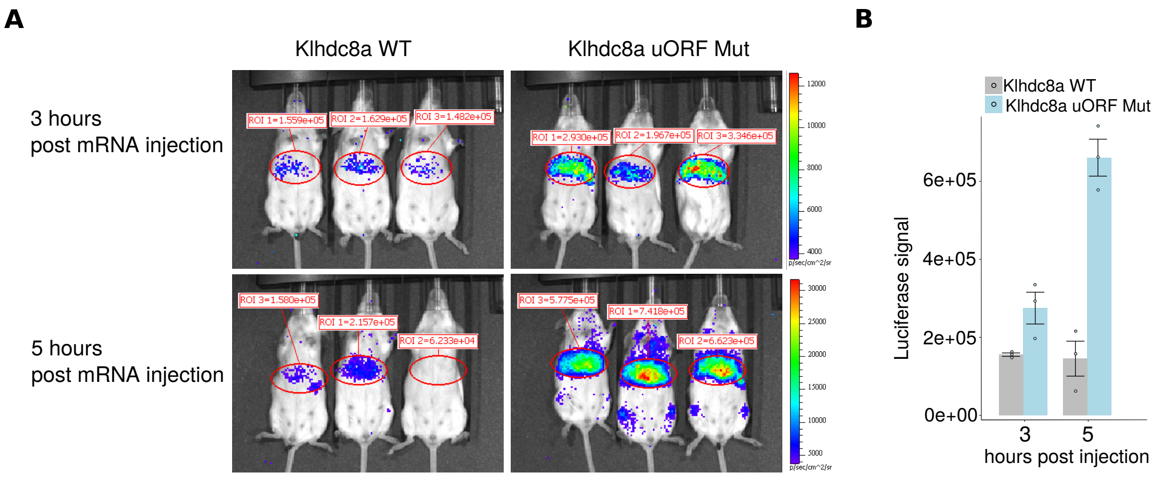
A
Klhdc8a WT
Klhdc8a uORF Mut
3 hours
post mRNA injection
5 hours
post mRNA injection
ROI 1=1.559e+05
ROI 2=1.629e+05
ROI 3=1.482e+05
ROI 1=2.930e+05
ROI 2=1.967e+05
ROI 3=3.346e+05
ROI 3=1.580e+05
ROI 1=2.157e+05
ROI 2=6.233e+04
ROI 3=5.775e+05
ROI 1=7.418e+05
ROI 2=6.623e+05
B
Klhdc8a WT
Klhdc8a uORF Mut
Luciferase signal
6e+05
4e+05
2e+05
0e+00
3
5
hours post injection

**Supplementary Figure 6. In vivo-bioluminescence imaging. A)** Synthetic mRNA with a WT or mutated *Klhdc8a* 5′ UTR and being fully substituted with N1-methyl pseudouridine (to avoid induction of type I interferon *in vivo*) was injected intravenously in mice using a formulation that allows expression of the *in vitro*-transcribed mRNA primarily in the liver. The luciferase activity was recorded *in vivo* 3 and 5 hours after the injection of RNA. Numbers indicate radiance in the region of interest (ROI). **B)** Quantification of the bioluminescence experiments in A. Error bars indicate mean $\pm$ sd.

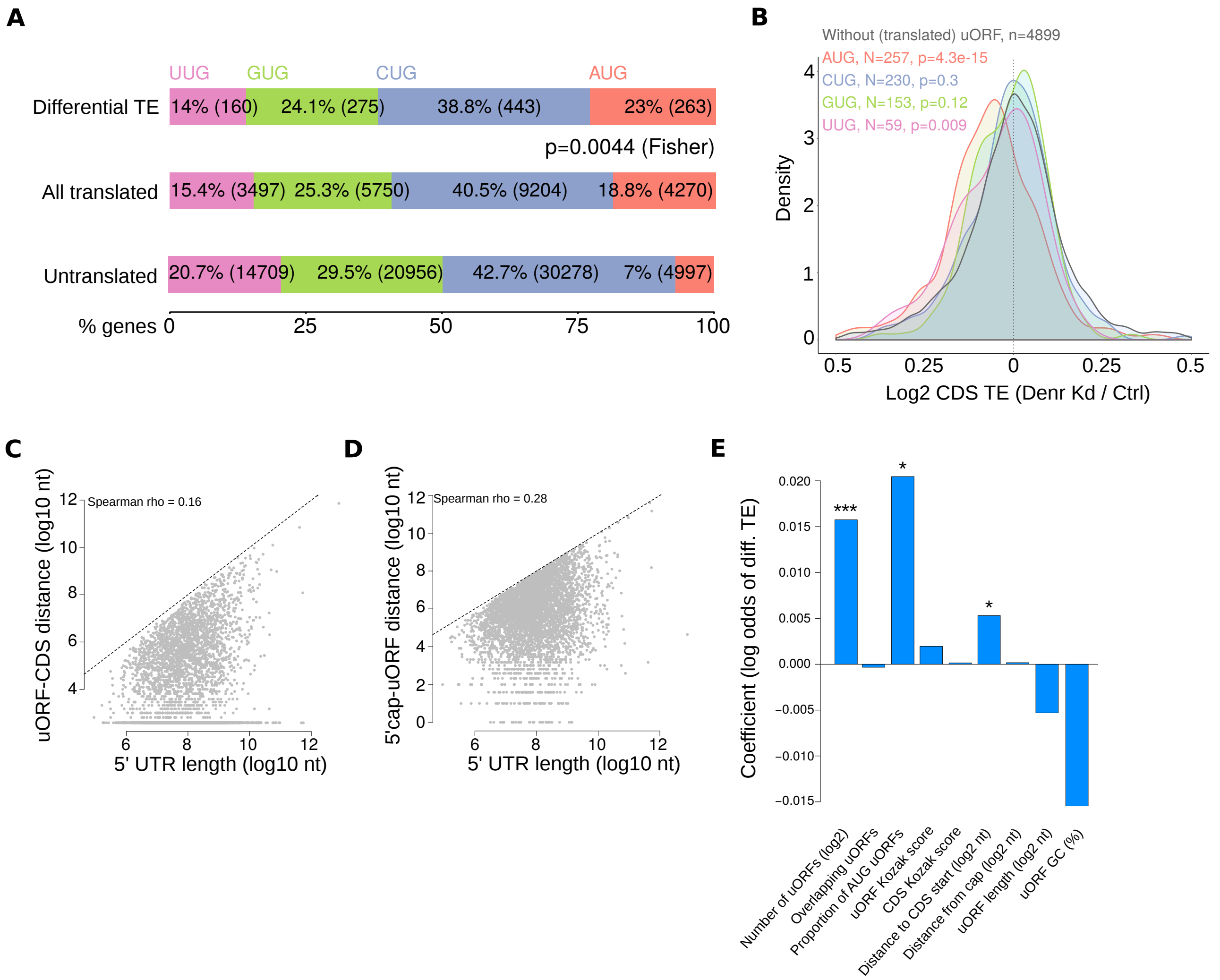
A
UUG
GUG
CUG
AUG
Differential TE
14% (160)
24.1% (275)
38.8% (443)
23% (263)
p=0.0044 (Fisher)
All translated
15.4% (3497)
25.3% (5750)
40.5% (9204)
18.8% (4270)
Untranslated
20.7% (14709)
29.5% (20956)
42.7% (30278)
7% (4997)
% genes
B
Without (translated) uORF, n=4899
AUG, N=257, p=4.3e-15
CUG, N=230, p=0.3
GUG, N=153, p=0.12
UUG, N=59, p=0.009
Density
Log2 CDS TE (Denr Kd / Ctrl)
C
Spearman rho = 0.16
uORF-CDS distance (log10 nt)
5' UTR length (log10 nt)
D
Spearman rho = 0.28
5'cap-uORF distance (log10 nt)
5' UTR length (log10 nt)
E
Coefficient (log odds of diff. TE)
Number of uORFs (log2)
Overlapping uORFs
Proportion of AUG uORFs
uORF Kozak score
CDS Kozak score
Distance to CDS start (log2 nt)
Distance from cap (log2 nt)
uORF length (log2 nt)
uORF GC (%)

**Supplementary Figure 7. Regression analyses of TE regulation by DENR. A)** Proportion of AUG, CUG, GUG, and UUG uORFs detected as translated, untranslated and among differential TE genes. DENR targets are enriched for translated AUG uORFs ($p = 0.0044$, Fisher test), indicating start codon specificity among DENR-regulated transcripts. **B)** Density plot of the CDS TE ratio (*Denr* Kd/control) for genes containing only AUG ($n = 257$), only CUG ($n = 230$), only GUG ($n = 153$), or only UUG ($n = 59$) uORFs. P-values indicate Wilcoxon test results for the difference between each distribution and that of genes without uORFs (or with untranslated uORFs) ($N = 4899$). Although the number of genes per category is low, this analysis shows that AUG-containing genes have the strongest CDS TE regulatory effect upon *Denr* knockdown. **C)** Scatter plot of $5'$ UTR length vs. the distance from the last uORF stop codon to the CDS start for all uORF-containing genes used for the regression model. No strong correlation was quantified between the two variables, ruling out that the correlation of the intercistronic distance with the magnitude of DENR regulation was due to longer $5'$ UTRs. **D)** Scatter plot of $5'$ UTR length vs. the distance from the $5'$ cap to the first uORF start codon for all uORF-containing genes used for the regression model. No strong correlation was quantified between the two variables, but correlation was higher than in (C). **E)** Coefficients for the logistic regression model, with "detected as differential TE, TRUE/FALSE" as binary response. Coefficients thus indicate the log odds change for being detected as differential TE (by Xtail) per unit change in the predictor. Same genes as for Figure 4 were used. Asterisks indicate coefficient estimates significance as: $p < 0.05$ (*), $p < 0.01$ (**), $p < 0.001$ (***).

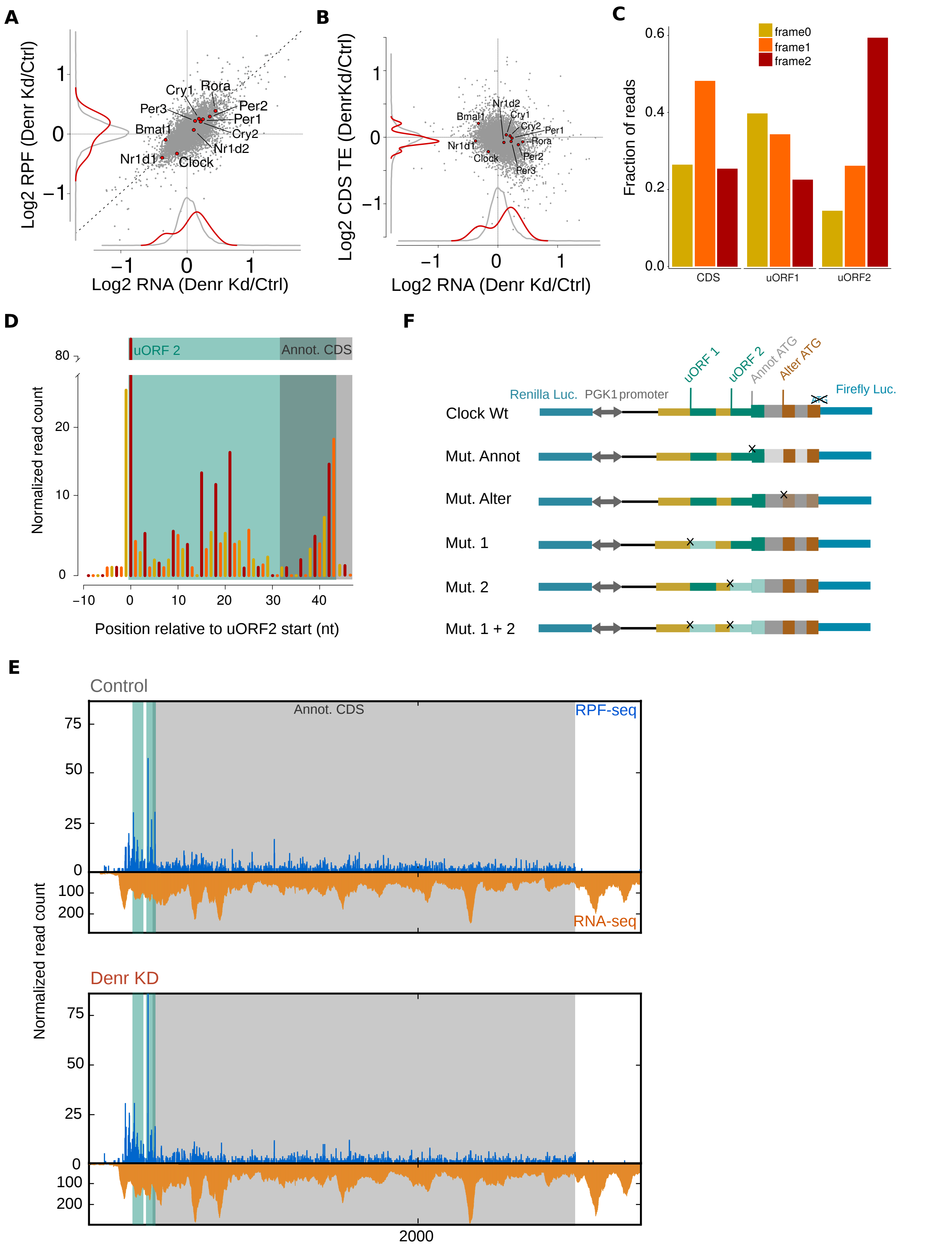
A
Log2 RPF (Denr Kd/Ctrl)
Log2 RNA (Denr Kd/Ctrl)
Cry1
Rora
Per3
Per2
Per1
Bmal1
Cry2
Nr1d2
Nr1d1
Clock
B
Log2 CDS TE (DenrKd/Ctrl)
Log2 RNA (Denr Kd/Ctrl)
C
Fraction of reads
frame0
frame1
frame2
CDS
uORF1
uORF2
D
uORF 2
Annot. CDS
Normalized read count
Position relative to uORF2 start (nt)
F
uORF 1
uORF 2
Annot ATG
Alter ATG
Renilla Luc.
PGK1 promoter
Firefly Luc.
Clock Wt
Mut. Annot
Mut. Alter
Mut. 1
Mut. 2
Mut. 1 + 2
E
Control
Annot. CDS
RPF-seq
RNA-seq
Normalized read count
Denr KD
2000

**Supplementary Figure 8. Clock is a DENR target. A)** Scatter plot of *Denr* Kd-to-control transcript levels (RNA-seq) vs. footprint levels (RPF-seq), with circadian core clock genes highlighted. *Clock* showed decreased translation without changes in mRNA abundance. **B)** Scatter plot of *Denr* Kd-to-control transcript levels (RNA-seq) vs. CDS translation efficiency, with circadian core clock genes highlighted. **C)** Frame preference of RPF-seq reads of *Clock* CDS, and both AUG uORFs. Frame preference within uORFs is similar to that on the CDS. **D)** Triplet periodicity within the uORF2. **E)** Distribution of normalised RPF-seq (blue) and RNA-seq (orange) reads along the *Clock* transcript, in control (top) and *Denr* Kd (bottom) cells. Green shaded area indicates the AUG-initiated uORFs; grey area corresponds to the annotated CDS. Only the first 500 nt of the 3′ UTR are shown for better visualisation. **F)** Schematic representation of the *Clock* wild-type and mutant constructs tested for effect on reporter synthesis and DENR dependence. The *Clock* reporters contain the whole 5′ UTR and the first 14 annotated amino acids of the CDS, in fusion with Firefly luciferase cDNA from which the start codon was removed. In addition, combinations of the mutated sites were cloned.

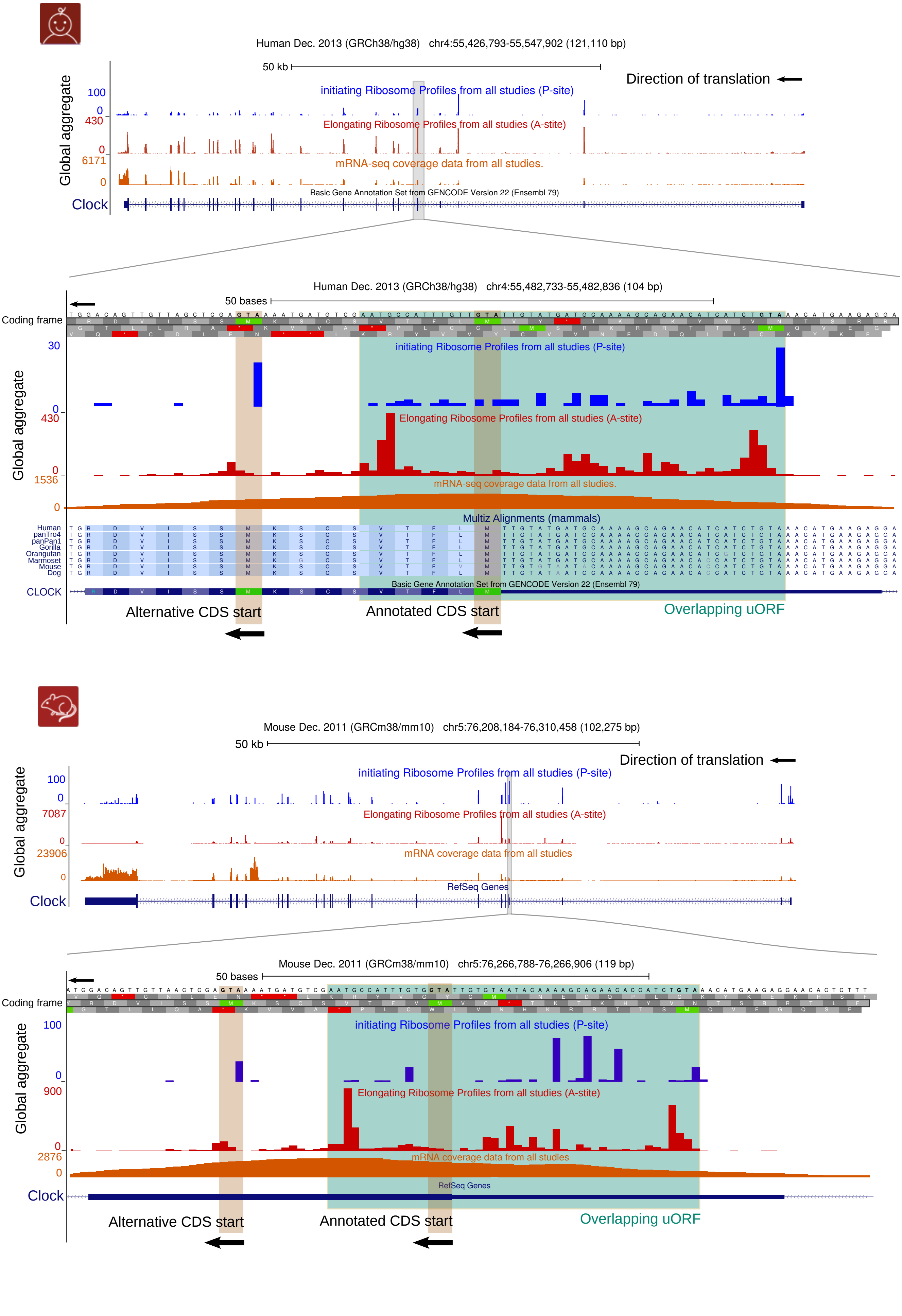

Human Dec. 2013 (GRCh38/hg38) chr4:55,426,793-55,547,902 (121,110 bp)
50 kb
Direction of translation
Global aggregate
100
0
initiating Ribosome Profiles from all studies (P-site)
430
0
Elongating Ribosome Profiles from all studies (A-site)
6171
0
mRNA-seq coverage data from all studies.
Basic Gene Annotation Set from GENCODE Version 22 (Ensembl 79)
Clock
Human Dec. 2013 (GRCh38/hg38) chr4:55,482,733-55,482,836 (104 bp)
50 bases
Coding frame
30
0
430
0
1536
0
Multiz Alignments (mammals)
Human
panTro4
panPan1
Gorilla
Orangutan
Marmoset
Mouse
Dog
CLOCK
Alternative CDS start
Annotated CDS start
Overlapping uORF
Mouse Dec. 2011 (GRCm38/mm10) chr5:76,208,184-76,310,458 (102,275 bp)
50 kb
Direction of translation
100
0
7087
0
23906
0
mRNA coverage data from all studies
RefSeq Genes
Clock
Mouse Dec. 2011 (GRCm38/mm10) chr5:76,266,788-76,266,906 (119 bp)
50 bases
Coding frame
100
0
900
0
2876
0
Alternative CDS start
Annotated CDS start
Overlapping uORF

**Supplementary Figure 9. Clock ribosome profiling data from published studies are consistent with alternative initiation from a downstream in-frame AUG.** All available data from initiating and elongating ribosome profiling studies, gathered from GWIPS-viz (46). An overview of the whole *Clock* transcript and a zoom over the CDS start is shown, where reads mapping on the overlapping uORF (over green background) and annotated and alternative CDS starts are visualised. Both human (top) and mouse (bottom) data are displayed. Initiating ribosomes can be seen at the start of the uORF and alternative CDS start. Elongating ribosome data (similar to our study) show good uORF coverage.

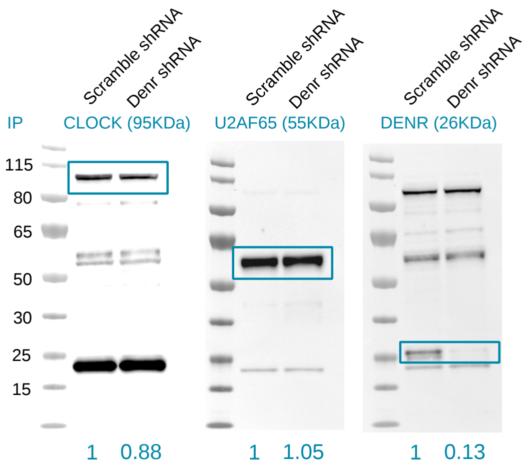
Scramble shRNA
Denr shRNA
Scramble shRNA
Denr shRNA
Scramble shRNA
Denr shRNA
IP
CLOCK (95KDa)
U2AF65 (55KDa)
DENR (26KDa)
115
80
65
50
30
25
15
1
0.88
1
1.05
1
0.13

**Supplementary Figure 10. Reduced CLOCK protein levels in Denr-depleted cells.** Left panel: Analysis of CLOCK levels in cells depleted of *Denr*. Middle panel: Same membrane probed against protein U2AF65, to asses equal loading. Right panel: Same membrane probed against DENR, indicating efficient knockdown. Numbers in blue indicate quantification of the signal using ImageJ, with Scramble shRNA-treated cells set to 1.
